# A unifying computational account of temporal context effects in language across the human cortex

**DOI:** 10.1101/2023.08.03.551886

**Authors:** Vy Ai Vo, Shailee Jain, Nicole Beckage, Hsiang-Yun Sherry Chien, Chiadika Obinwa, Alexander G. Huth

**Affiliations:** Intel Labs, Intel Corporation; Hillsboro, OR 97124, USA; Department of Computer Science, The University of Texas at Austin; Austin, TX 78712, USA; Department of Psychological and Brain Sciences, Johns Hopkins University; Baltimore, MD 21218, USA; Department of Electrical & Computer Engineering, The University of Texas at Austin; Austin, TX 78712, USA; Department of Neuroscience, The University of Texas at Austin; Austin, TX 78712, USA

**Keywords:** fMRI, human cortex, timescales, natural language/speech

## Abstract

Deep learning advances have revolutionized computational modeling approaches in neuroscience. However, their black-box nature makes it challenging to use deep learning models to discover new insights about brain function. Focusing on human language processing, we propose a new framework to improve the quality and interpretability of the inferences we make from deep learning-based models. First, we add interpretable components to a deep language model and use it to build a predictive encoding model. Then, we use the model’s predictive abilities to simulate brain responses to controlled stimuli from published experiments. We find that our model, based on a multi-timescale recurrent neural network, captures many previously reported temporal context effects in human cortex. Its failure to capture other effects also highlights important gaps in current language models. Finally, we use this new framework to generate model-based evidence that supports the proposal that different linguistic features are represented at different timescales across cortex.

## Introduction

The rise of artificial neural networks (ANNs) has created a revolution in cognitive and systems neuroscience. Despite their many differences with biological brains, ANNs trained to perform tasks such as image classification, spatial navigation, and language modeling have been shown to capture a surprising array of behavioral and neurophysiological phenomena^1,2^. ANNs have been used to create encoding models (EMs) that predict brain responses to naturalistic visual, auditory, and language stimuli independent of the neuro-imaging/recording technology^3-8^, providing additional evidence of their alignment to biology. Some have asserted that neural language models (LMs) are the most effective computational models we have of human language processing^9,10^. Autoregressive LMs are trained on the task of predicting the next word in a stream of text, and have been shown to learn a variety of useful representations that transfer to other language tasks^11,12^. However, the "black box" nature of ANNs has made it difficult to accept these as computational models of the brain. Ideally, computational models should undergo iterative refinement to reveal the useful and necessary principles of neural processing that give rise to experimentally observed phenomena.

A comprehensive model of human language processing would capture a variety of experimental effects, such as the effects of temporal context. Prior work has shown that language-responsive regions in the human cerebral cortex appear to be organized across a hierarchy of processing timescales that capture varying amounts of context. While some regions respond to short timescale information (single words), others integrate over longer timescales (sentences or paragraphs)^13,14^. Neural LMs have also been shown to effectively process information at many different temporal scales^15-18^, but there is typically no explicit temporal processing hierarchy in these models^19^. Therefore it is not known whether neural LMs can capture the variety of temporal context effects observed in humans. Here we demonstrate two possible solutions to the problem of testing ANNs as computational models of the brain. These solutions will enable further work to advance research into both neural LMs and into the cognitive neuroscience of natural language processing.

Our first approach to this problem is to replace "black box" ANNs with more interpretable models that contain components which can be linked to known theoretical constructs. We use a *multi-timescale recurrent neural network (MT-RNN) LM* where each unit in the model processes information at a fixed, known timescale^17,20^ (**Figure 1A**). The unit timescales follow a distribution that allows the MT-RNN to model the power law decay typically seen in natural language and other hierarchically structured data^21,22^ (**Figure 1B**). We then build a multi-timescale encoding model (MT-EM) (**Figure 1C**) from the MT-RNN. These encoding models are trained using neuroimaging data in which seven participants each listened to over six hours of naturalistic narratives from *The Moth Radio Hour* while whole brain blood oxygen level dependent (BOLD) responses were recorded with fMRI^23^. The added interpretability of this model reveals which timescale units are important for predicting responses in each brain region, providing a direct estimate of representational timescales in the brain^20^ (**Figure 1D-E**). Our second approach is to conduct *in silico* experiments that test the ability of these interpretable EMs to capture different aspects of language processing in the brain. Here, we use the MT-EMs to simulate brain responses under stimulus manipulations used in past neuroimaging studies of temporal context effects^14,24,25^ (**Figure 1F**). If the MT-EM makes similar predictions as the past studies, it would suggest that the MT-RNN is a good computational model of temporal processing in the brain. If it does not match past predictions, it could reveal shortcomings of the MT-RNN or the assumptions of the modeling approach.

**Figure 1.**
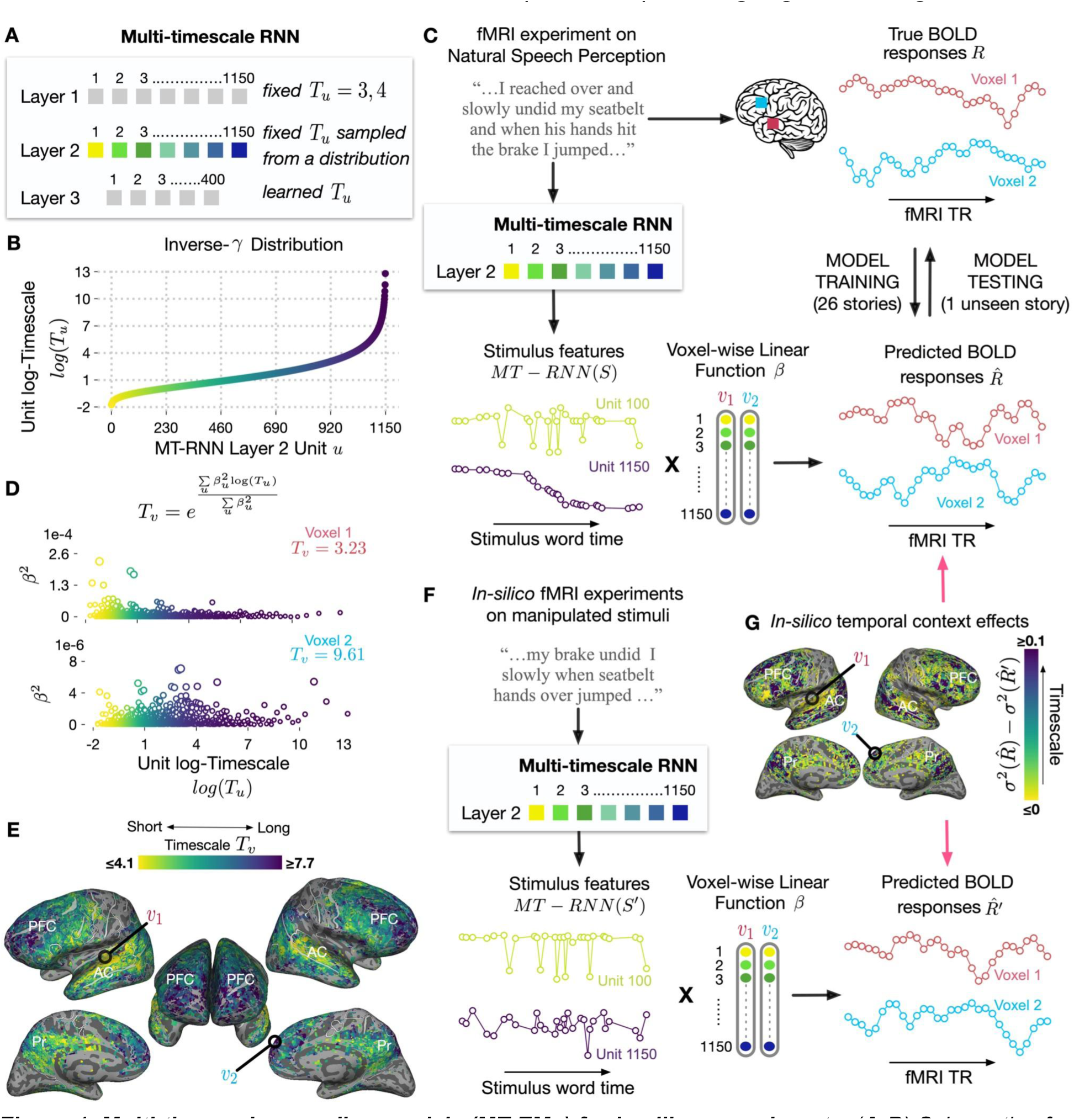
Multi-timescale encoding models (MT-EMs) for in silico experiments. (**A-B**) Schematic of 3-layer multi-timescale RNN (MT-RNN) language model (LM). In layer 1, the timescale T_u_ of each MT-RNN unit u was fixed at 3 or 4 words. In layer 2, T_u_ followed an inverse gamma distribution sampled at 1150 points. In layer 3, T_u_ was freely learnable. (**C**) To build a model that predicts fMRI responses, we extracted encoding model features from all 1150 MT-RNN layer 2 units after they had processed each word of the speech stimuli. Linear regression was used to learn a mapping between the features and the measured BOLD response for each voxel, β_v_. The performance of the multi-timescale encoding models (MT-EM) was evaluated by comparing the true and predicted responses on held-out data. (**D**) Distribution of MT-EM regression weights, β, across MT-RNN Layer 2 units for two different voxels. Although many units have non-zero weights for each voxel, short timescale units contribute more towards predicting voxel 1’s responses while longer timescales contribute the most for voxel 2. By taking a weighted average across unit timescales, we can estimate the timescale T_v_ of each voxel.(**E**) T_v_ estimates for statistically T_v_ significant cortical voxels in one participant (all in **Supp. Figure S2**). Results corroborate the temporal hierarchy reported in prior work^14,20^, with short-timescales in AC and longer timescales in PFC and Pr. Following T_u_, the T_v_ units are in words (see **Methods** for details). (**F**) In silico evaluation framework testing MT-EM’s ability to capture temporal context effects reported in previous studies. Each original study compared cortical BOLD responses to an intact story S and its temporally manipulated version S’. We extracted MT-RNN features for S and S’ and simulated BOLD responses to them given our previously learned voxel weights. (**G**) Example in silico results that preserve single-voxel, single-participant detail. Each in silico experiment depends on a metric that compares the responses to the true and manipulated stimuli, 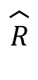 and 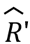. PFC: Prefrontal cortex; AC: Auditory cortex; Pr: Precuneus.

The key contributions of our work are empirical, meta-scientific, and theoretical. Our empirical results show that multi-timescale encoding models successfully simulate previously reported effects^14,24,25^ on how linguistic information is constructed over time. This suggests a computational isomorphism between the MT-RNN and the human brain. However, we also find that it does not reproduce a report on how contextual information is *forgotten*^25^, pointing to the need for novel ANN architectures that can capture this phenomenon. Despite this shortcoming, the MT-EMs are more comprehensive computational models than previous efforts^25-27^ because they both account for the linguistic content of the stimuli at different timescales and directly predict brain responses in individual participants and voxels. This makes the MT-EM a comprehensive, unified computational account of temporal context effects in human cortex during natural language processing. Our second, meta-scientific contribution is the *in silico* experimentation framework that leverages the strengths of ANNs and predictive capabilities of EMs to advance questions in cognitive neuroscience^28^. These *in silico* experiments provide challenging generalization tests for encoding models that go beyond measuring prediction performance on held-out data from the same task. *In silico* experiments can both evaluate brain-ANN correspondence and suggest new adaptations that may even improve performance on machine learning tasks like language modeling^28^. Lastly, we leverage the interpretability of the LM to explore the relationship between timescales and linguistic content in the MT-RNN^17,29^. Through this analysis, we make our third, theoretical contribution which proposes that different linguistic features may be processed in different timescale-specific regions of cortex. In sum, these results open up several new avenues of investigation that leverage the strengths of deep learning and experimental neuroscience.

## Results

### Multi-timescale Recurrent Neural Networks (MT-RNNs) for language encoding

To build an interpretable neural language model, we constructed the multi-timescale recurrent neural network (MT-RNN) by explicitly defining a timescale *T_u_* for each unit *u* in the first two layers of the model (**Figure 1A**). This time constant controls the exponential rate of memory decay: the strength of a memory will decay by a factor of *e* after *T_u_* words^15^. In the first layer, *T_u_* was fixed to remember information over either 3 or 4 words. In the second layer, the 1150 different unit timescales followed an inverse gamma distribution (**Figure 1B**) in which units remember between three and thousands of words. In the last layer, unit timescales were left unrestricted to aid MT-RNN in the language modeling task. Like all auto-regressive LMs, MT-RNN takes in one word at a time from a training document and learns to output a probability distribution that predicts the next word. It was trained on English language Wikipedia^30^ documents, and fine-tuned on narrative stories similar to the fMRI stimuli (**Methods**). While MT-RNN is marginally worse at language modeling than contemporaneous transformer architectures, its advantage lies in the improved timescale interpretability.

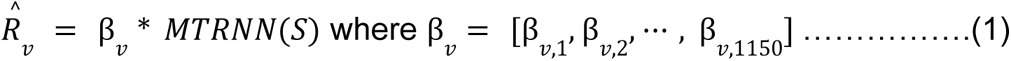

Next, we used this interpretable LM to build voxelwise encoding models. The trained MT-RNN was first used to process each narrative story in the fMRI stimulus set. For each word in the story we extracted 1150-D feature vectors from layer 2 units, resulting in representations that span a broad set of timescales. Then cross-validated ridge regression was used to learn a 1150-D linear mapping, β_*v*_, from the MT-RNN features of the story to the elicited BOLD *T_v_* response, *R_v_*, of each cortical voxel *v* in each participant (**Figure 1C**; **Equation 1**; **Methods**). We measured prediction performance of this multi-timescale encoding model (MT-EM) for each voxel *v* by correlating between true, *R_v_*, and predicted, 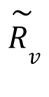, responses for held-out data not used during the estimation of β_*v*_. To test whether prediction performance was significantly greater than chance, we built leave-one-training-story-out encoding models and ran blockwise permutation tests on the prediction performance of the left-out stories (*p < 0. 001*, FDR corrected^31^). We only report results for significantly predicted voxels in each participant (significant voxel count *µ = 18, 378*; *σ = 6, 402*).

In all participants, MT-EM significantly predicted responses across the temporal, parietal and frontal lobes (**Supp. Figure S1**), replicating earlier results with the same model^20^. This illustrates that a distributed network of regions across cortex robustly responds to the multi-timescale linguistic information captured by the MT-RNN.

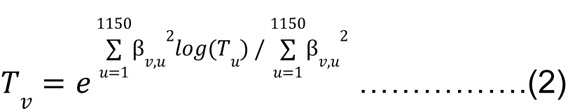

The estimated encoding model weights for each voxel, β_*v*_, tell us how much the 1150 different units-each with a known timescale-contributes to the voxel response (**Figure 1D**). We used these weights to create a model-based estimate of each voxel’s average processing timescale, *T_v_*, in every participant by computing the center-of-mass across unit timescales *T_u_* weighted by the corresponding model weights β_*v*_ (**Equation 2**). Our assumption is not that individual voxels operate at a single timescale, but that their apparent timescale is a combination of many underlying neurons. We discuss different ways to interpret the units of the model-based timescale estimate *T_v_* in the **Methods**. But like other timescale estimates, larger *T_v_* values indicate that information is retained over longer temporal windows.

**Figure 1E** shows the cortical distribution of model-based timescales *T_v_* in an example participant (all participants in **Supp. Figure S2**). Overall, *T_v_* appears hierarchically organized across the cortical surface, with short timescales appearing in early stages of speech processing (superior temporal lobe, posterior frontal lobe) and long timescales in prefrontal cortex and angular gyrus. These findings replicate earlier encoding model work^20,32^ and broadly corroborate the organization proposed by earlier studies^14,20,33-37^.

Although MT-EM has good prediction performance overall, this evaluation is limited to naturalistic speech perception. It does not demonstrate that MT-EM is a *comprehensive* model of language processing which can capture other known temporal context effects. To evaluate this, we developed an *in silico* framework that tested MT-EM’s ability to simulate experimental effects from three published *in vivo* studies^14,24,25^. First, the experimental stimuli were manipulated as described in the original study (for example, the order of words was shuffled). The trained MT-EMs were then used to predict BOLD responses to these manipulated stimuli (**Figure 1F**; **Equation 1**). This is a difficult generalization test for the MT-RNN and MT-EM, as they had not previously been exposed to manipulated stimuli. Finally, we devised a series of analyses to test the impact of the experimental manipulation on each voxel as a function of its timescale (**Figure 1G**). While each analysis preserved the intent of the original experiment, we made modifications to enable single-participant, single-voxel resolution of the effects (**Methods**).

### Temporal scrambling experiment

We simulated Lerner et al.’s influential study on scrambling natural stories to uncover a temporal hierarchy across cortex^14^. The authors recorded BOLD responses while participants listened to a narrative story or to temporarily manipulated versions of that story, obtained by playing it backwards, scrambling the order of words (Duration per word: µ = 0. 7s; σ = 0. 5), sentences (µ = 7. 7s; σ = 3. 5; ∼9 words), or paragraphs (µ = 38. 1s; σ = 17. 6; ∼55 words). They showed that some cortical regions were unaffected by scrambling over longer durations, indicative of short processing timescales, while other long-timescale regions were significantly affected by all scramble conditions. To simulate this experiment, we scrambled word chunks in each story of the fMRI stimulus set 100 different times. This was done at the granularity of single words, 9 words (sentence-level), or 55 words (paragraph-level). We report similar results for stories manually segmented into sentences and paragraphs (**Supp. Figure S5**).

First we examined whether the timescales of MT-RNN units affected their responses to each scramble condition. We fed each word of the intact (*S*) or scrambled story (*S*’) to the MT-RNN and extracted responses from layer 2 units. To compute a correlation between responses to *S* and *S’*, we re-ordered each unit’s response to match the original word order (**Figure 2A**). A short-timescale unit should be indifferent to any scrambling (**Figure 2B**, left), while a unit processing sentences should be affected by word scrambling but not sentence or paragraph (**Figure 2B**, right). We found that word-level scrambling affected all MT-RNN units, although the drop in correlation was larger for long-timescale units (**Figure 2C**). Sentence-level scrambling had little effect on short-timescale units, but substantially affected units with timescales over the average sentence length (*T_u_ > 9*). Paragraph-level scrambling greatly affected units with*T_u_ > 55*, the average paragraph length. To further quantify this effect, we built a linear model to predict each unit’s *T_u_* from the z-transformed correlation values of each scrambling condition. All of the conditions were significant predictors (coefficients for word: 0.478; sentence: -1.163; paragraph: -1.3104) for a model *R^2^* of 0.759. Overall, these results illustrate that MT-RNN does capture the effects tested by the scrambling paradigm.

**Figure 2.**
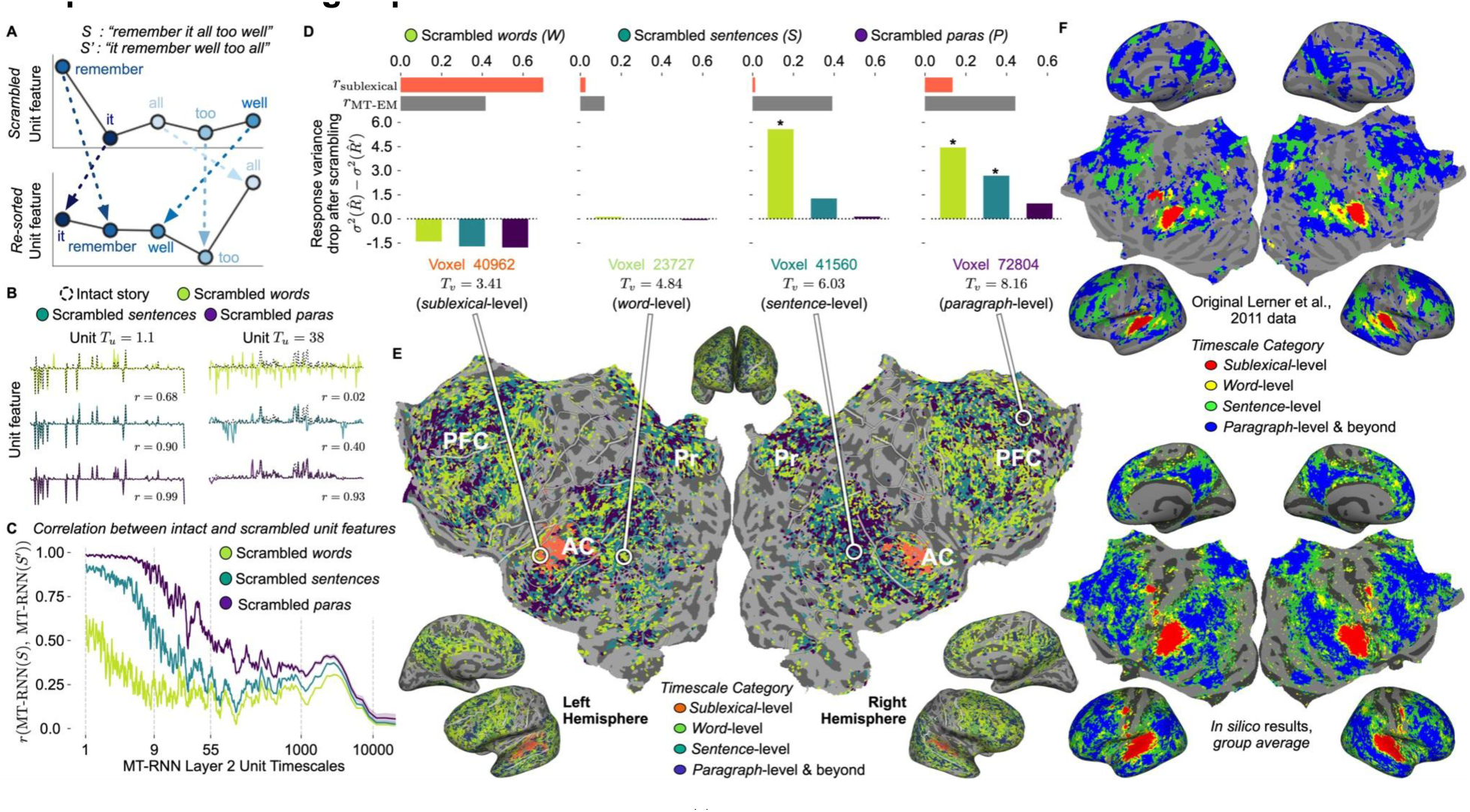
In silico replication of Lerner et al., 2011^14^, which presented subjects with a story scrambled at the word-, sentence-, or paragraph-level. (**A**) MT-RNN features extracted from a story scrambled at the word-level. To make a direct comparison with the intact story features, the scrambled features were re-sorted. (**B**) Effect of story scrambling on example short (left) and long timescale unit (right) representations. Word-level scrambling caused the largest difference between the intact (dashed line) and scrambled (solid color) responses. However, the short-timescale unit was less affected by word-level scrambling than the long-timescale unit. The longer timescale unit was affected by word- and sentence-level scrambling, but not paragraph-level. (**C**) Correlation between intact and scrambled MT-RNN features for all layer 2 units, where r = 1.0 indicates no difference in the MT-RNN unit’s response to the intact and scrambled stories. Scrambling every word of the story (green) resulted in the largest drop in correlation. However, short-timescale units (left) were less affected by word-scrambling. Longer timescale units that integrated over a paragraph (∼T_u_ = 55) showed a larger drop in correlation to sentence- and word-level scrambling than paragraph-level. (**D**) Categorical timescale assignments based on voxel responses to different scramble conditions. "Sublexical-level" voxels were better predicted by a sub-lexical encoding model (**Methods**) than MT-EM. Word-level voxels showed no significant change in response across all scramble conditions (i.e. unaffected by word scrambling and beyond). Sentence-level voxels only showed a significant change to word scrambling. Voxels with a significant change to at least word and sentence scrambling conditions were labeled "paragraph-level & beyond". (**E**) Distribution of categorical voxel timescales across cortex in an example participant. (**F**) A comparison between the group-averaged results of our in silico experiment (N=7) and the original study (N=11). The group-averaged maps smooth over variability seen in individual subject maps (**Supp. Figure S3**). Abbreviations follow Figure 1.

Next, we estimated cortical timescales by simulating each voxel’s responses to scrambled stories and comparing it to the intact version. In the original study, this was done by spatially co-registering participants into a shared anatomical space and measuring the inter-subject correlation (ISC) for each scrambling condition. Here, we measured the variance of the predicted response for each condition in each voxel of each participant. Since response variance monotonically varies with ISC (**Methods**), this measure allowed us to conduct a similar analysis but preserve individual differences and voxel-level resolution. We could not directly simulate voxel responses to the stories played backwards, since MT-RNN is a word-level model. Instead, we built a sub-lexical encoding model that captures sub-word properties like phonemic content (**Methods**) and labeled any voxels better predicted by this model than MT-EM as "sub-lexical". The remaining voxels were sorted into timescale buckets (*word*, *sentence*, and *paragraph*) based on changes in response variance across scrambling conditions (**Figure 2D-E**). Averaging results across participants revealed that the timescale buckets estimated with our *in silico* framework matched the temporal hierarchy across cortex reported by the original study (**Figure 2F**). However, the single-participant, single-voxel estimates allowed us to capture more detailed variation in regions like prefrontal cortex, precuneus and angular gyrus (**Figure 2E**).

We then tested whether this experiment-based timescale estimate correlated with our model-based *T_v_* metric. The estimated timescale buckets were significantly but weakly correlated with the *T_v_* estimates of MT-EM in each participant (r = 0.055 ± 0.016 SEM; prevalence test across participants p < .001). As discussed in a later section (**A unifying, model-based metric of temporal context effects**), the strength of this correlation is mediated by the performance of the encoding model.

In **Supp. Figure S4**, we show MT-EM results from two correlation-based metrics instead of response variance. Although each metric broadly produces similar results as **Figure 2E**, we demonstrate how underlying assumptions in a metric or statistical testing procedure can bias observed results.

Overall, this conceptual replication provides evidence that MT-EM captures temporal context effects tested by the scrambling paradigm. The interpretability of MT-EM and the ability to link the temporal scrambling experiment with model-based *T_v_* highlight an important difference between these results and previous work that used transformer-based LMs to simulate this experiment^38^.

### Narrative divergence experiment

Next, we performed an *in silico* experiment closely following the manipulation from Yeshurun et al., 2017^24^ (**Figure 3A**). There, the authors presented participants with two versions of a narrative story that only differed in about ½ of the words. These words were semantically similar but led to subtle changes in the meaning of each story. The study found that voxels in long timescale regions showed a greater distance between BOLD response timecourses to the two story variants. This suggested that these regions accumulated differences between the stories over a longer span of time. To simulate this experiment, we presented the two story versions used in that study to MT-RNN and extracted features for each word in the stories.

**Figure 3.**
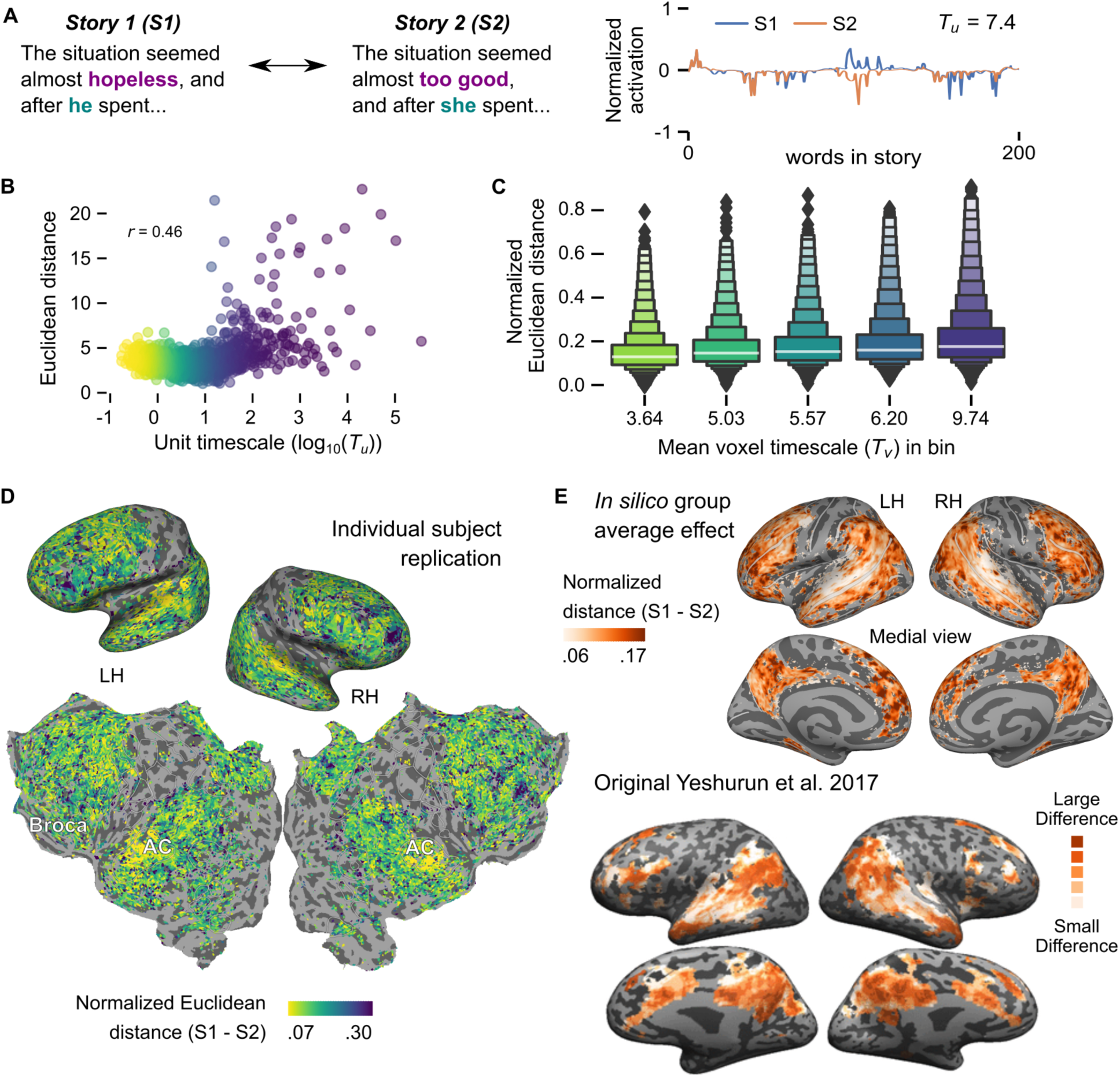
The in silico replication of Yeshurun et al. 2017^24^, which replaced some of the words in a story to change its overall meaning. They found that the BOLD responses to Story 1 (S1) and Story 2 (S2) diverged over time, with long timescale regions showing a larger difference between the two stories. (**A**) An example of the words replaced between S1 and S2 (left), and the response of a single MT-RNN unit to the first 200 words in either story (right). (**B**) Replicating this experiment on the MT-RNN shows a correlation between unit timescale and the normalized Euclidean distance between story responses. (**C**) In silico replication effect across participants. Plot shows the average correlation between average voxel timescale T_v_ (in 5 evenly sized bins) and normalized Euclidean distance between stories. (**D**) Normalized Euclidean distance between stories for an example participant. This value is smaller in short-timescale regions, such as AC, and greater in longer-timescale regions, such as Broca’s area. (**E**) A comparison between group-averaged results from our in silico experiment (N=7) and the original study (N=18). Abbreviations follow Figure 1.

We first investigated the effect of this manipulation on MT-RNN units by computing the Euclidean distance between each unit’s normalized feature timecourse for the two stories (**Methods**). Following the original study, we hypothesized that the distance should scale with the assigned timescale of each unit. Indeed, the correlation between log-transformed unit timescales and their Euclidean distances was significant (r = 0.46; p < 0.001 permutation test; FDR q < .05) (**Figure 3B**). This suggests that activations of MT-RNN units diverge between the two stories if the units have longer timescales, much like cortical voxels.The MT-RNN is thus able to capture the temporal context effect tested by this paradigm.

Next, we used the MT-RNN features for each story to simulate cortical BOLD responses with MT-EM. To enable comparisons across voxels, we normalized the simulated voxel responses by the magnitude of the encoding model weights. This controls for the fact that ridge regression can shrink the magnitude of the predicted responses differently in each voxel (**Methods**). We then computed the Euclidean distance between each voxel’s normalized response timecourse for the two stories, and scaled this metric between 0 and 1 for each participant. We found that Euclidean distances were significantly correlated with model-based *T_v_* in each participant (r = 0.19 ± 0.004 SEM; prevalence test across participants p < .001; **Figure 3C**). Furthermore, the pattern of results across the cortical surface broadly matched the original study-while voxels in short timescale regions like primary auditory cortex had small distances, long timescale regions like inferior parietal cortex had much larger distances (**Figure 3D**). Much like the temporal scrambling experiment, we found more detailed variations in individual participant maps compared to group-averaged maps (**Figure 3E**).

The previous *in silico* experiment manipulated the coherent temporal structure of language inputs to observe how different brain regions process information at many timescales. This experiment uses stimuli that only differ in semantic content while maintaining a similar temporal structure. Overall these results illustrate that the MT-EM also captures temporal context effects that depend purely on semantic differences.

### Constructing and forgetting context experiment

Finally, we examined a study by Chien and Honey that measured the rate at which different cortical regions construct or forget prior context in a narrative^25^. Using data from the paragraph scrambling condition in Lerner et al., 2011^14^, the authors measured how context representations were constructed by computing a timepoint-by-timepoint correlation between BOLD responses to the same paragraph when the preceding context varied (**Figure 4A**). To measure how context was forgotten, they computed the correlation between responses to different paragraphs preceded by the same context. The study found that the rate of context construction scaled with the purported temporal hierarchy across cortex. In contrast, the rate of forgetting across a paragraph was similar across cortical regions.

**Figure 4.**
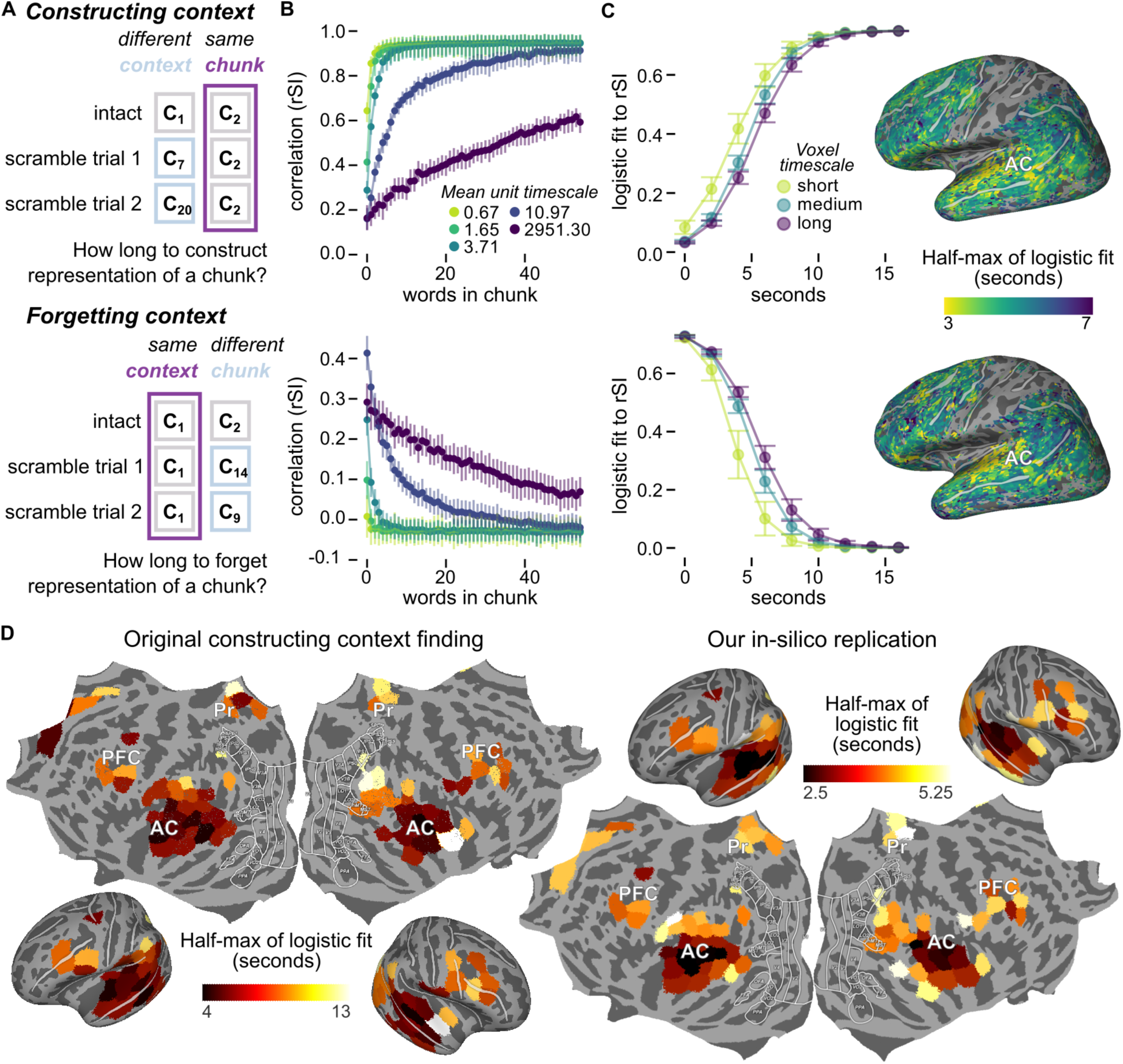
The in silico replication of Chien et al. 2020, which analyzed the time it takes to construct or forget prior context in natural language. For (**A**)-(**C**), constructing context effects are shown in the top row, and forgetting context in the bottom row. (**A**) An illustration of the experimental manipulation which measured how long it takes to construct or forget context. Using the same scramble trials from our Lerner et al. 2011 in silico experiment, we analyze chunks based on whether they are preceded or followed by the same chunk. (**B**) We extracted MT-RNN feature timecourses for the intact and scrambled stories, and computed the correlation between them for every word in a 55-word chunk (rSI). A high correlation shows that the unit’s feature is very similar between the intact and scrambled stories, i.e. context is no longer affecting the unit’s activity. MT-RNN units with a short timescale T_u_ construct and forget context very quickly (light green), whereas units with a long T_u_ (dark purple) are still influenced by contexts longer than 55 words. Error bars are 95% bootstrap CIs. (**C**) We repeated this analysis on the predicted fMRI timecourses generated from the MT-EM. To simplify the display of the results, we fit a logistic function to the correlation curves. Left: the averaged fits for voxels that are in the bottom 5% (short), middle 5% (medium), or top 5% (long) of voxel timescales T_v_ (right). Short timescale voxels both construct and forget context more quickly (light green). Right inset: logistic fit half-maximums shown for an example participant. While our data replicates the original constructing context finding, it does not replicate the original forgetting effect. Error bars are SEM across participants. (**D**) A direct comparison between the original constructing context effect (left, N=31) and our in silico replication effect averaged across participants (right, N=7). We are able to show these on the same cortical map because we had full access to the original data. Abbreviations follow Figure 1.

To investigate the context construction and forgetting behavior of MT-RNN, we first divided units into evenly-binned timescale groups. For each timescale bin, we conducted a similar timepoint-by-timepoint correlation analysis as the original work. We reused the scrambled 55-word, paragraph-level chunks from the *in silico* Lerner experiment to generate 100 trials that matched the context construction or context forgetting conditions (**Figure 4A**). We show the average correlation within each timescale bin in **Figure 4B**. We found that the rate of context construction across a paragraph was slower in longer timescale units, much like the published report. However, unlike the published cortical data, the rate of forgetting across the paragraph also varied with unit timescale *T_u_* instead of remaining constant.

Next, we performed a similar analysis using the predicted BOLD responses in the paragraph scrambling condition. In the original study, the authors spatially co-registered participants into a common anatomical space and obtained average timecourses with variability across participants. To maintain resolution at a single-participant, single-voxel level, we modified this procedure and obtained average timecourses with variability across scramble trials (**Methods**). Such an approach would be difficult if not impossible to do *in vivo* since it would involve a substantial amount of data collection per participant. Following the original study, we then fit a logistic function to each voxel’s correlation values across timepoints in a paragraph for both construction and forgetting. We found that the construction time, estimated as the half-maximum of the logistic curve, was slower for voxels with longer model-based *T_v_* estimates, aligning with prior findings (**Figure 4C-D**). There were more detailed variations in the individual participant maps than the group-averaged results (**Figure 4E**), similar to the *in silico* experiments above.

However, we found that the forgetting time also varied with *T_v_*, diverging from the original findings (**Figure 4D**). The correlation between average constructing and forgetting times across voxels was significant within each of the 7 participants (r = 0.568 ± 0.006 SEM; prevalence test p < 0.001). To compare with the original study, we also computed this correlation by averaging data within different regions-of-interest across participants (r = 0.944; p < 0.001 permutation test; FDR corrected) and in the group-averaged maps (r = 0.840; p < 0.001 permutation test; FDR corrected). Again these results were in contrast to the original study, which did not find a significant correlation between constructing and forgetting times across ROIs.

One possible explanation for the discrepancy in forgetting results could be that the MT-RNN language model is incapable of forgetting information quickly, or has not been trained to do so. To investigate this hypothesis, we evaluated MT-RNN performance on the language modeling task in 3 different conditions: each paragraph was either preceded by the correct context in the story, the wrong context due to paragraph scrambling, or no context at all (**Supp. Figure S6**). We found that language modeling performance was better when the model had no access to context than when the context was wrong/shuffled. This provides evidence that MT-RNN is incapable of forgetting old, irrelevant context, even when forgetting would improve its ability to predict the next word. The inability of the MT-RNN to forget information may be due to common choices in the preprocessing of discourse and context markers in LM training data (**Methods**). Alternatively, the MT-RNN may not learn human-like forgetting behavior until it provides a large performance benefit on the next-word prediction task. Improving the forgetting behavior of the MT-RNN may require changes to its forgetting mechanisms, to its training procedure, to the training data, to the objective function, or all of the above. These would all increase the strength of the training signal to forget information, and may improve the overall performance of the MT-RNN or LMs generally on other downstream language tasks.

### A unifying, model-based metric of temporal context effects

Through the *in silico* framework, we have shown that the MT-EM can account for many different temporal context effects across cortex. However, it remains unclear whether these timescale-related metrics are all capturing the same information, and how different biases in the modeling procedure or experimental paradigm might affect the results we observe. One strength of the *in silico* framework is that it enables us to directly compare across experimental paradigms and examine different aspects of our modeling assumptions. To address these points, we evaluated the relationship between each replicated effect and the model-based timescale estimate, *T_v_*. We also computed correlations with SNR (**Methods**) and MT-EM prediction performance. This was motivated by two observations: first, that metrics like response variance and intersubject correlation are confounded by the quality of the fMRI signal (**Methods; Temporal scrambling experiment**), and second, that the scale of the predicted fMRI signal varies with the regularization parameter that is estimated separately for each voxel (**Methods; Narrative divergence experiment**). In **Figure 5** we show the pairwise correlation between *T_v_* and each experiment-based timescale metric: timescale categories from temporal scrambling (*Lerner RV*), Euclidean distance between diverging narratives (*Yeshurun d_norm_*), and the time it took to construct a context representation (*Chien T_c_*). For each pair of metrics, we first computed the mean correlation within participant, and then averaged across participants. Since all of our timescale-related metrics depend on the MT-EM, we also compared them to a model-free timescale estimate derived from the coherence of the actual BOLD signal in each voxel (*T_mf_*), inspired by prior work on timescales^39^. This provides a test of whether our *in silico* estimates are capturing timescale information that is independent of the quality and assumptions of the modeling procedure itself.

**Figure 5.**
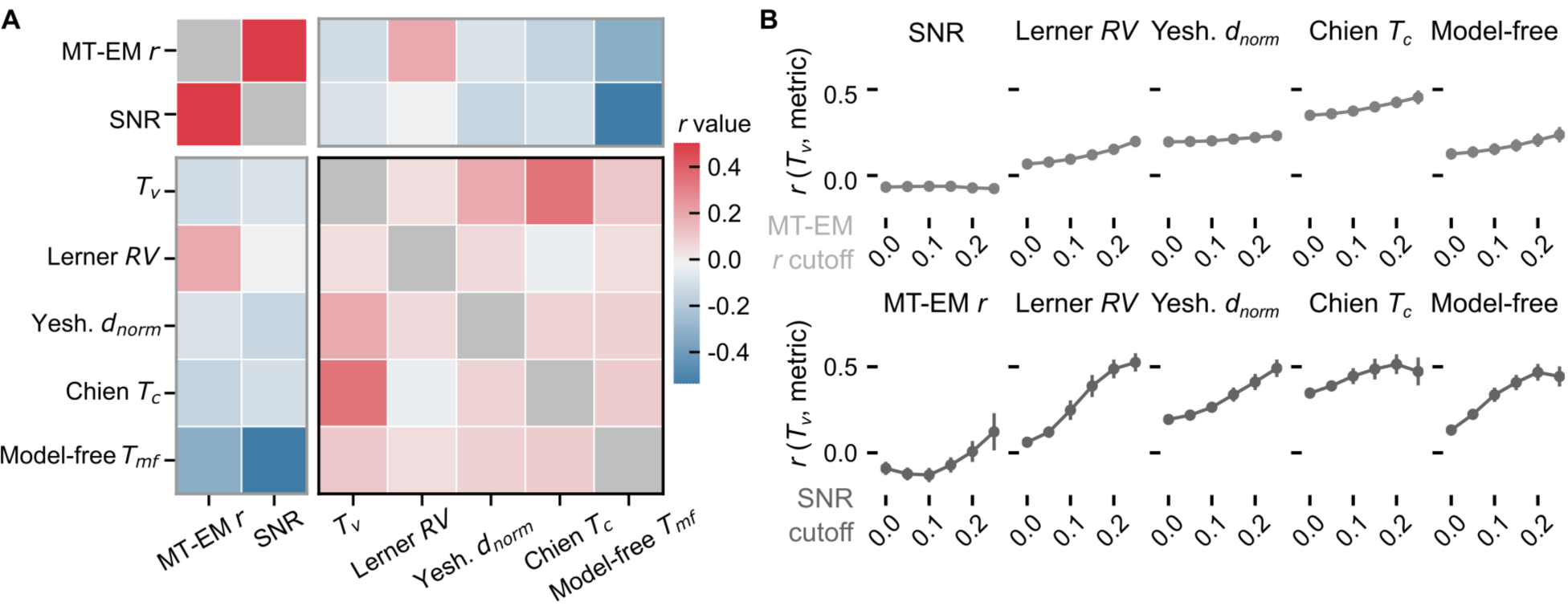
The relationship between MT-EM derived timescale T_v_, the in silico experiment metrics that successfully replicate the original reports, a model-free timescale metric T_mf_ based on spectral information, and metrics related to the quality of the fMRI signal. Abbreviations: Lerner RV is the timescale category based on fMRI response variance (Figure 2E).Yeshurun d_norm_ is the normalized Euclidean distance between the stories (Figure 3C). Chien T_C_ is the half-max of the logistic fit for the constructing context condition (Figure 4C). MT-EM r is the measure of encoding model performance (**Fig 1C**), and SNR is the signal-to-noise ratio. (**A**) Cross-correlation matrix of timescale metrics, averaged within participant and then across participants. (**B**) Correlations re-computed only for voxels that survive a threshold on encoding model performance (top) or SNR (bottom). Error bars show standard error over participants. The T_v_ metric more reliably predicts the in silico timescale metrics as signal quality and encoding model performance become more reliable. As the thresholds increase, the correlation between timescale T_v_ and the in silico experiment metrics generally increases. The effect is weaker when thresholding MT-EM r for the Yeshurun metric since we normalize d_norm_ by the magnitude of the MT-EM weights. Error bars are SEM across participants.

Overall, the model-based estimate, *T_v_*, best encompassed other timescale metrics, with an average correlation of 0.197 ± 0.049 SEM across experiment-based metrics (**Figure 5A**). *T_v_* was not affected by SNR (r = -0.068 ± 0.059 SEM) or MT-EM prediction performance (r = -0.090 ± 0.043 SEM). In contrast, *Lerner RV* was much more sensitive to MT-EM prediction performance (r = 0.197 ± 0.024 SEM). We hypothesized that this confound might explain the weak correlation we observed between *Lerner RV* and *T_v_*. To further examine how the MT-EM or recorded fMRI signal quality might affect the temporal context metrics, we evaluated each metric’s relationship with *T_v_* at different performance thresholds. The correlation for the model-free metric and most of the timescale metrics increased monotonically with the threshold (**Figure 5B**; top panel). Only *Yeshurun d_norm_* was flat across performance thresholds. This is because we normalized the predicted BOLD signal here by the magnitude of the MT-EM weights (**Methods**). When we thresholded voxels by SNR, we also found that the correlation between *T_v_* and other metrics increased monotonically with the threshold (**Figure 5B**; bottom panel). These analyses suggest that the correlations between metrics could improve with increased data quality and better encoding models, and provide further evidence that these metrics are all capturing similar covariance driven by a voxel’s processing timescale.

To follow the original studies, we also re-analyzed these relationships after aligning all participants into a common anatomical space (**Supp. Figure S8**). Since collapsing data across participants reduces noise, correlations were higher between most timescale metrics. However, the relationship of *T_v_* with SNR (r = 0.023), and MT-EM prediction performance (r = -0.026) remained weak. This shows that *T_v_* can also capture group-level cortical variability in the temporal context effects, although MT-EM uniquely facilitates single voxel, single-participant resolution. One key advantage of *T_v_* is that estimating it does not require additional data collection or simulation beyond the natural speech perception experiment. This may be advantageous over using unnatural stimuli, like scrambled stories, that could elicit compensatory effects *in vivo*. Given these results, we propose that the multi-timescale encoding model is a comprehensive and unifying computational account of many temporal context effects in human cortex during language processing.

### Probing the timescale of functional content in MT-RNN

How does the timescale of a cortical brain region interact with its selectivity for different linguistic features? Earlier studies have shown that the cortical areas we analyze by timescale are selective for different linguistic features such as semantic concepts, syntactic operations or discourse processing^41-47^. However, these studies did not simultaneously examine the timescale of a region. Since neural network language models capture many linguistic features at different scales^29,48-50^, we can use the MT-RNN to estimate both cortical timescale and linguistic selectivity^51-54^. In the final analysis, we examined the relationship between timescales and different linguistic features in MT-RNN to understand the functional roles of different timescales during language processing.

First, we extracted MT-RNN features for different pieces of text and divided them into five non-overlapping buckets based on their timescale-short phrases (*T_u_* ≤ 1; 0-3 words), sentences (1 ≤ *T_u_* ≤ 9; ∼8 words), long sentences (9 ≤ *T_u_* ≤ 20; ∼18 words), paragraphs (20 < *T_u_* ≤ 70; ∼28 words), and multiple paragraphs (*T_u_* > 70; more than 70 words) (see **Supp. Figure S9** for other splits). Then we built linear classifiers that used each set of MT-RNN features to predict four different linguistic annotations of the text at different scales: part-of-speech (word), named entity type (word), entailment or contradiction (sentence), and topic (paragraphs). This approach is based on popular interpretive tools for ANNs^48,55-58^. We compared the classifier performance of different timescale buckets against a classifier that used the entire feature space.

Classifier performance on word-level tasks (part-of-speech and named entity) was worse for longer timescale units (**Figure 6**; errorbars represent 95% CIs). Moreover, the shortest timescale bucket performed comparably with the full feature space, suggesting that the shortest timescale units in MT-RNN were enough to encode individual word features. In contrast, topic information was primarily encoded in long-timescale units, with near-chance performance for shorter timescales. Unlike the other tasks, the sentence entailment task had comparable performance across all timescales. This could be because the other timescale buckets relied on confounding heuristics like word matching to solve the task^40^. Overall, our results show that MT-RNN units encode different linguistic information based on their timescales. Coupled with our earlier findings, these results provide evidence that the linguistic function of each voxel may differ depending on its processing timescale. This is contrary to proposals that language processing operates over the same linguistic ‘grain size’^59^.

**Figure 6.**
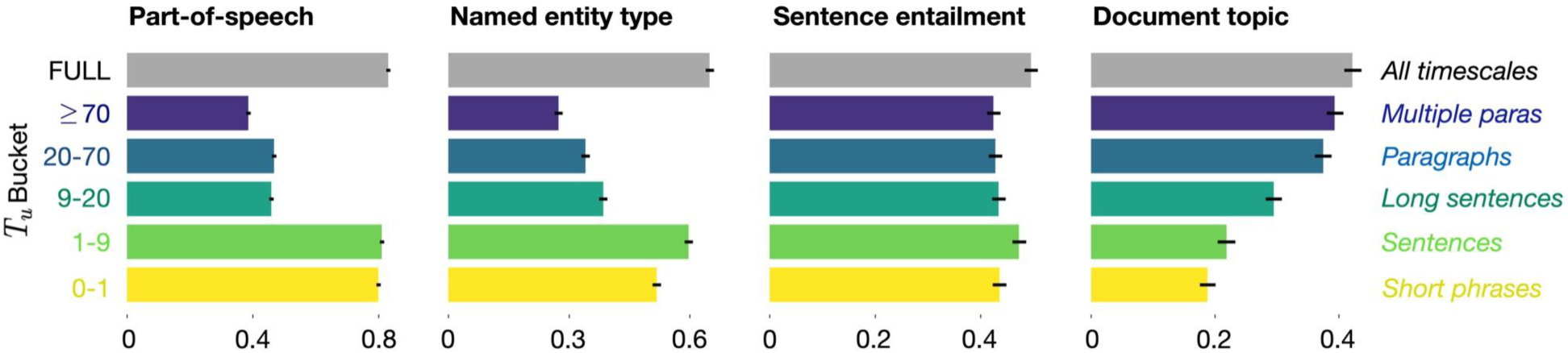
To understand the types of linguistic information encoded by MT-RNN units at different timescales, linear classification probes were built for four different NLP tasks operating over different linguistic scales. Left to right: part-of-speech (word); named entity type (word): classifying named entities like person or place; sentence entailment (sentence): textual entailment task determining if two sentences are contradictory, neutral or one entails the other; document topic (paragraph). Linear classifiers were either trained using the full 1150-D MT-RNN feature space or a subset of the features in specific timescale buckets. As expected, short timescale units predicted part-of-speech tags almost as well as the full model. Named entities were best predicted by a slightly longer timescale bucket (1-9) and topic information was best predicted by the longest timescale bucket (>=70). Despite being a sentence-level task, all the buckets performed similarly for sentence entailment (possibly due to short timescale confounds in the task^40^.) Error bars represent 95% bootstrap CIs.

## Discussion

Here we showed that an interpretable computational model of language processing, the MT-RNN, can be fit to brain data collected while humans listened to natural language to produce a multi-timescale encoding model (MT-EM). We used the MT-EM to conceptually replicate temporal context effects reported in three earlier fMRI studies. We also probed the MT-RNN and found that different types of linguistic information were represented at different timescales: short-timescale units contained information about features such as part-of-speech, while longer timescale units contained more information about high-level features such as topic. This provides new support for the idea that different timescale-specific regions of cortex process different linguistic features.

Our study contributes to a growing body of work on ANN-based encoding models^9,20,52,60-62^ and provides an example of *in silico* experimentation with these models. This approach has several advantages (see Jain et al.^28^ for thorough discussion). First, we can effectively model ecologically relevant tasks like natural language perception^63^ and test how these models generalize to controlled stimuli in theoretically interpretable experimental designs. This allows us to leverage the respective strengths of different experimental approaches. Second, *in silico* experimentation allows us to easily analyze how experimenter choices may affect any observed results to improve reproducibility. Another conceptual replication of the Lerner et al. study, for example, chose a different metric to summarize the differences across conditions^38^. We show a similar correlation-based metric in **Supp. Figure S4**. In both cases, these metrics result in a high prevalence of ‘word-level’ regions, illustrating that the choice of metric can affect scientific inferences. Future *in silico* work should investigate other potential sources of bias, such as different ANNs, random seeds, training data diversity, or even different distributions of model parameters such as *T_u_*. This is essential to understand the robustness and generalizability of the results. Ramifications of experimenter choices are also facilitated by sharing code and data, as we do for this work^64^.

Another important future direction is building better, more comprehensive models of temporal context effects in the brain. Our results showed that MT-RNN could simulate temporal effects that involved constructing context, but could not simulate forgetting behavior^25^. A recent report found that reservoir networks could capture many temporal context effects, including forgetting^27^. However, this work assumed that a separate mechanism had already learned how to represent each word in the stimuli. Future efforts should focus on building models that can overcome the drawbacks of both these approaches. One might look to popular transformer architectures to model temporal context effects, which can outperform the MT-RNN even after controlling the number of parameters^17^. Yet methods to control timescales in transformers are limited^65^, and indirect methods to estimate cortical timescales in individual transformer layers^60,61^ are not as reliable^20^ as directly controlling model timescale. By contrast, the MT-RNN is fairly biologically plausible. Each MT-RNN unit’s activity exhibits exponential decay, and the distribution of MT-RNN units allows it to approximate scale-invariant power-law behavior^17^ observed in natural sequential data^6^. This distribution is a plausible approximation of single neuron time constant distributions reported across varying tasks and brain regions^66-68^. Future *in silico* experimentation will need to balance task performance against biological plausibility to address neuroscience questions about temporal context effects.

Unlike previous studies of timescales during language processing^14,24,25,69^, our estimation procedure provides single-participant, single-voxel resolution. This can facilitate future translational work between data modalities (e.g. electrophysiology and fMRI^66^). Our results also provide single-voxel level evidence for heterogeneous timescales within a cortical region, as has been reported for other tasks and species^68^. This heterogeneity may provide important clues to how different neural mechanisms drive both intrinsic and functional timescales^67,68^. A within-participant comparison of intrinsic timescales measured from resting state fMRI versus functional timescales measured during different tasks could provide further mechanistic insight.

There is a growing body of literature on how functional demands modulate apparent timescale^68-71^. In the domain of language, functional timescales are consistent between spoken and written language despite different rates of information transmission^32^. Timescales are also adaptable to the rate of speech as long as it is intelligible^69^. This flexibility suggests that perhaps the linguistic content of the stimuli is more important than the intrinsic timescale. Here we used linear probes as a preliminary effort to understand what linguistic content may be represented at different scales. Recent work explored this in more depth by examining the relationship between semantic selectivity and integration timescales across the cortical surface (Jain et al., *in prep*).

In agreement with prior work^35,46^, they found that semantic selectivity is a major principle of functional organization across cortex during language processing in comparison to timescales. However, this is only the beginning of an effort to understand how different types of linguistic information may be computed at different temporal scales. Future work that explores both the semantic content and temporal dynamics across different tasks will be critical to improve our understanding of how different timescales enable us to process real-world stimuli.

## Methods

### fMRI experiment on natural language perception

We used our publicly released fMRI dataset on natural language perception^72^. This dataset comprises seven human participants each listening to over five hours of naturalistic, narrative English-language stories.

Data collection was broken up into 6 different scanning sessions, the first session involving the anatomical scan and localizers, and each successive session consisting of 5 or 6 stories. One of the 27 stories (∼10 minutes long) was reserved for testing the encoding models and was played five times for each participant-once during each scanning session. The five responses were then averaged across the repeats.

#### Naturalistic narrative dataset

The fMRI stimuli comprised 27 English-language narrative stories from *The Moth Radio Hour* (∼10-15 minutes long each). These rich, complex stimuli were highly representative of language that humans encounter on a daily basis and spanned many different semantic concepts spread across different timescales. In each story, a single speaker told an autobiographical story without reading from a prepared speech. The stories contained 57,899 words in total out of which 5,206 were unique. Each story was transcribed and the transcript was aligned to the audio to find the exact time each word was spoken.

#### Stimulus Preprocessing

Each story was manually transcribed by one listener. Certain sounds (for example, laughter and breathing) were also marked to improve the accuracy of the automated alignment. The audio of each story was then downsampled to 11kHz and the Penn Phonetics Lab Forced Aligner^73^ (P2FA). This software was used to automatically align the audio to the transcript. Praat^74^ was then used to check and correct each aligned transcript manually.

#### Stimulus preparation and presentation

During the fMRI experiment, story stimuli were played over Sensimetrics S14 in-ear piezoelectric headphones. The audio for each story was filtered to correct for frequency response and phase errors induced by the headphones using calibration data provided by sensimetrics and custom python code (https://github.com/alexhuth/sensimetrics_filter). All stimuli were played at 44.1 kHz using the pygame library in Python (https://www.pygame.org/news).

#### Participants

Data were collected from three female and four male human participants: UT-S01 (author SJ, female, age 24), UT-S02 (author AGH, male, age 34), UT-S03 (male, age 22), UT-S05 (female, age 24), UT-S06 (female, age 24), UT-S07 (male, age 24), UT-S08 (male, age 24). We follow the same participant ID mapping here as the original data release^72^. All participants were healthy and had normal hearing, and normal or corrected-to-normal vision. The experimental protocol was approved by the Institutional Review Board at the University of Texas at Austin. Written informed consent was obtained from all participants. To stabilize head motion during scanning sessions participants wore a personalized head case that precisely fit the shape of each participant’s head.

#### Acquisition parameters

MRI data were collected on a 3T Siemens Skyra scanner at the UT Austin Biomedical Imaging Center using a 64-channel Siemens volume coil. Functional scans were collected using a gradient echo EPI sequence with repetition time (TR) = 2.00 s, echo time (TE) = 30.8 ms, flip angle = 71°, multi-band factor (simultaneous multi-slice) = 2, voxel size = 2.6mm x 2.6mm x 2.6mm (slice thickness = 2.6mm), matrix size = 84×84, and field of view = 220 mm.

Anatomical data for all participants except UT-S-02 were collected using a T1-weighted multi-echo MP-RAGE sequence on the same 3T scanner with voxel size = 1mm x 1mm x 1mm following the Freesurfer morphometry protocol. Anatomical data for participant UT-S-02 were collected on a 3T Siemens TIM Trio scanner at the UC Berkeley Brain Imaging Center using a 32-channel Siemens volume coil using the same sequence.

#### Processing fMRI data

All functional data were motion corrected using the FMRIB Linear Image Registration Tool (FLIRT) from FSL 5.0. FLIRT was used to align all data to a template that was made from the average across the first functional run in the first story session for each participant. These automatic alignments were manually checked for accuracy.

Low frequency voxel response drift was identified using a 2nd order Savitzky-Golay filter with a 120 second window and then subtracted from the signal. To avoid onset artifacts and poor detrending performance near each end of the scan, responses were trimmed by removing 20 seconds (10 volumes) at the beginning and end of each scan, which removed the 10-second silent period and the first and last 10 seconds of each story. The mean response for each voxel was subtracted and the remaining response was scaled to have unit variance.

#### Flatmap Reconstruction

Cortical surface meshes were generated from the T1-weighted anatomical scans using FreeSurfer software^75^. Before surface reconstruction, anatomical surface segmentations were hand-checked and corrected. Blender was used to remove the corpus callosum and make relaxation cuts for flattening. Functional images were aligned to the cortical surface using boundary based registration (BBR) implemented in FSL. These alignments were manually checked for accuracy and adjustments were made as necessary.

Flat maps were created by projecting the values for each voxel onto the cortical surface using the "nearest" scheme in the pycortex software package^76^. This projection finds the location of each pixel in the flat map in 3D space and assigns that pixel the associated value. This was done by finding the pixel locations halfway between the pial and white matter surfaces.

#### Localizers

Known regions of interest (ROIs) were localized separately in each participant. Three different tasks were used to define ROIs; a visual category localizer, an auditory cortex localizer, and a motor localizer.

For the visual category localizer, data was collected in six 4.5 minute scans consisting of 16 blocks of 16 seconds each. During each block 20 images of either places, faces, bodies, household objects, or spatially scrambled objects were displayed. Participants were asked to press a button every time an image was presented twice in a row. The corresponding ROIs defined in the cerebral cortex with this localizer were the fusiform face area (FFA), occipital face area (OFA), extrastriate body area (EBA), parahippocampal place area (PPA), and occipital place area (OPA).

The motor localizer data was collected during 2 identical 10-minute scans. The participant was cued to perform six different tasks in a random order in 20 second blocks. The cues were ‘hand’, ‘foot’, ‘mouth’, ‘speak’, saccade, and ‘rest’ presented as a word at the center of the screen, except for the saccade cue which was presented as an array of dots. For the ‘hand’ cue, participants were instructed to make small finger-drumming movements for the entirety of the cue display. For the ‘foot’ cue, participants were instructed to make small foot and toe movements. For the ‘mouth’ cue, participants were instructed to make small vocalizations that were nonsense syllables such as *balabalabala*. For the ‘speak’ cue, participants were instructed to self-generate a narrative without vocalization. For the saccade cue, participants were instructed to make frequent saccades across the display screen for the duration of the task.

Weight maps for the motor areas were used to define primary motor and somatosensory areas for the hands, feet, and mouth; supplemental motor areas for the hands and feet, secondary somatosensory areas for the hands, feet, and mouth, and the ventral premotor hand area. The weight map for the saccade responses was used to define the frontal eye fields and intraparietal sulcus visual areas. The weight map for speech was used to define Broca’s area and the superior ventral premotor (sPMv) speech area (Chang et al. 2011).

Auditory cortex localizer data was collected in one 10 minute scan. The participant listened to 10 repeats of a 1-minute auditory stimulus containing 20 seconds of music (*Arcade Fire*), speech (Ira Glass, *This American Life*) and natural sound (a babbling brook). To determine whether a voxel was responsive to auditory stimulus, the repeatability of the voxel response across the 10 repeats was calculated using an *F-*statistic. This map was used to define the auditory cortex (AC).

The precuneus and prefrontal cortex (PFC) ROIs were defined by the Freesurfer parcellation for each participant. The PFC ROI was a combination of the superior frontal, rostral middle frontal, frontal pole, and caudal middle frontal parcellations.

### Multi-timescale Recurrent Neural Network (MT-RNN)

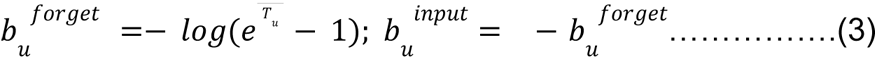

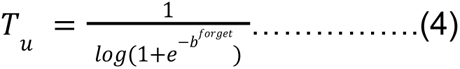

The multi-timescale recurrent neural network (MT-RNN) is an interpretable 3-layer long short-term memory (LSTM)^77^ network. In traditional LSTMs, each unit *u* has an associated *forget* gate which controls the rate at which *u* remembers past information and an *input* gate which controls the rate at which *u* integrates new information. To make the LSTM more interpretable, we made *u* operate at a specific timescale *T_u_* by fixing the bias factors of both these gates, *b_u_^forget^* and *b_u_^input^*, as specified in **Equation 3**. Since LSTM units generally exhibit exponential decay of information^15^, the unit timescale *T_u_* is simply the exponential time constant that specifies how much time is required for information in the unit to decay by *1Ie* (assuming the natural logarithm in **Equations 3-4**).

Following Mahto et al.^17^, we fixed the timescales of all 1150 units in layer 1 of MT-RNN such that they integrated over 3 or 4 words. In layer 2, the timescales of all 1150 units followed an inverse gamma distribution (**Figure 1A-B**). This initialization allowed the summed activity of all units to mimic an overall power law decay, which has been observed in natural language and other hierarchically structured data^21,22^. As layer 2 received inputs from layer 1, the effective timescale of each *u* in layer was *T_u_ + 3. 5*, where 3.5 is the estimated average timescale of layer 1 units. In reality, each layer 2 unit may receive an unevenly weighted combination of inputs from layer 1, so the timescale contribution of layer 1 would not exactly equal 3.5. Despite this we can interpret layer 2 units as integrating over about 3 words to thousands of words. The 400 units in layer 3 had freely varying timescales (i.e. learnable by backpropagation) so the MT-RNN could perform well on the task it was trained on-autoregressive language modeling (LM). In the LM task, the ANN has to learn to predict the next word in a sequence, *w_i_*, from all the past words, *w*_[1,*i*-1]_. Through this, the ANN can learn to capture many different types of linguistic information from low-level syntax to high-level discourse information^48-50,56,58,78-81^. While it is hard to interpret the timescale at which different linguistic information is captured in traditional LMs, the added interpretability of MT-RNN allows us to explore this directly.

We followed standard machine learning practices to pre-train the MT-RNN on common language modeling datasets. Details not described here can be found in the published paper on the MT-RNN^17^. The training data consisted of the WikiText-2 dataset^82^, which comprises ∼2 million words in the training set and ∼33,000 unique words in its vocabulary. Note that pre-processing included the removal of punctuation and line breaks, which are markers that could indicate the start of a new context. As with other LMs, the input to the MT-RNN was a learnable embedding layer which represented each unique word in the vocabulary as a 400-D vector. Any word outside the vocabulary was labeled with a special "<UNK>" token that was initialized to a zero embedding. Unlike most LMs, we maintained the sequential order of each document when training the MT-RNN LM. This is possible because we pass the hidden state from the previous context window to the next context window (i.e. "stateful" RNN training).

Since the naturalistic narrative podcasts we used in the fMRI experiment were stylistically different from the training data, we then fine-tuned MT-RNN to do language modeling on other stories from The Moth Radio Hour, TED Talks and Modern Love. This fine-tuning dataset had ∼940,000 words in total. We re-trained the embedding and output layers to use a vocabulary of ∼13,800 unique words that encompassed the fMRI stimuli. For all the analyses shown here, we froze the model after fine-tuning so its weights were no longer trainable.

The MT-RNN LM was trained and fine-tuned using a single GeForce GTX TITAN X GPU with 64GB CPU RAM. All code was written in pytorch^83^. Following prior work^82^, we used stochastic gradient descent (SGD) and subsequently, non-monotonically triggered average SGD for optimization. Additional details on the model and training can be found in Jain et al.^20^ and Mahto et al.^17^ (which has accompanying publicly available code).

### Multi-timescale fMRI encoding models for natural language perception

With their ability to capture many different types of linguistic information, neural LMs make very good predictive models of the brain. A growing body of research has demonstrated this across different presentation modalities (written or spoken), neuro-imaging/recording techniques (fMRI, MEG, ECoG etc.) and parts of the brain (cerebral cortex or cerebellum)^4,9,60-62,84-86^. Following this literature, we extracted the interpretable hidden state of layer 2 in MT-RNN to build multi-timescale encoding models (MT-EM) of the fMRI blood-oxygen-level-dependent (BOLD) signal during natural language perception. To extract these hidden states, we froze the fine-tuned model so its internal parameters could not be changed and then passed the word transcript of each fMRI stimulus into the model. We extracted a 1150-D multi-timescale feature representation for every word in the stimulus.

Since the BOLD signal was acquired at 0.5Hz, we had to down-sample the word-level features prior to modeling. In our prior work^20^, we showed that traditional temporal down-sampling techniques used in fMRI encoding models do not work well for long timescale features. Since these techniques assume that each word occurs as an infinitesimal spike, they can track features that rapidly change across words (like the part-of-speech) but cannot faithfully down-sample low frequency information (like the narrative’s topic). To this end, that work proposed an interpolation scheme based on radial basis function kernels whose widths were proportional to the timescale of each unit. We use the same downsampling mechanism here. Since this method accounts for the duration of each word, it could also be used for compressed/dilated speech^69^.

Once the MT-RNN features were downsampled to the rate of the fMRI stimuli, we used a finite impulse response (FIR) model with 4 delays to approximate the haemodynamic response function. We then trained an L2-regularized ridge regression model for each individual voxel in each participant. We refer to this as the multi-timescale encoding model (MT-EM). The MT-EM used the MT-RNN features to predict the BOLD activity elicited by listening to the narrative stories. Of the 27 stories, 26 were used to train the MT-EM. These stories were also used to find the voxel’s optimal ridge coefficient by repeating the regression fitting procedure 50 times. Each time a random sample of 5000 TRs (125 blocks of 40 consecutive TRs) from the training set was held-out from training and used to evaluate the performance of the model across different ridge coefficients. The best coefficient was chosen based on the average performance across all 50 repeats^46,87^.

To evaluate the prediction performance of each voxel *v*’s MT-EM, we first generated predictions, 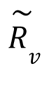, for the single held-out story not used during model estimation. Then, we computed the linear correlation *r_v_ ^test^* between 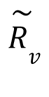 and true BOLD responses to the story, *R_v_*.

#### Statistical significance tests for encoding model performance

To evaluate whether encoding model performance was statistically significant, we used a leave-one-training-story-out fitting procedure on the training data. Specifically, we fit 26 separate encoding models by leaving out each of the 26 stories individually and generated predictions for each held-out story. Then we computed the linear correlation, *r_v_^loo^*, between the concatenated true responses for all held-out stories and the concatenated predicted responses. The statistical significance of *r_v_^loo^* was established with a permutation test. Since BOLD activity is heavily autocorrelated in time, the predicted responses were first divided into blocksof 10 consecutive TRs. Then an empirical null distribution was computed by randomly permuting the blocks 1000 times and recomputing the correlation each time. The original correlation, *r_v_^loo^*, was evaluated against this null distribution to get a p-value. To correct for multiple comparisons across voxels, we did False Detection Rate (FDR) correction (*q < 0. 001*)^31^. More details on the encoding model fitting procedure and significance testing can be found in our prior work^20^.

Since all the temporal context studies we simulate here operate in a group-averaged space, we also conducted statistical significance tests at the group-level. To do this, we first projected *r_v_^loo^* and its 1000 permuted versions into the Freesurfer fsaverage space. Then we computed p-values and did FDR correction to find the significant fsaverage voxels within each participant. Finally we computed the proportion of participants that showed a significant effect for each voxel. Using a bayesian procedure^88^, we estimate the prevalence of this effect. The group-averaged encoding model performance for significant voxels is shown in **Supp. Figure S1**.

#### Sub-lexical encoding models

The MT-RNN only produces features at a lexical level, which limits the sensitivity of the MT-EM to sublexical information. To identify cortical areas that are sensitive to sublexical information, we concatenated four different sublexical features and built encoding models with them. The first two features were the rate of words and phonemes in each stimulus story. The third feature was a 1-hot encoding space that comprised 44 dimensions, one for each phoneme in American English as defined by the CMU Pronouncing Dictionary^89^ and a few non-speech sounds like laughter or clapping. The last feature space was the place of articulation and voicing of each phoneme. We followed the same procedure as in MT-EM to establish statistical significance at the individual participant and group level.

#### Signal-to-noise ratio of cortical BOLD activity

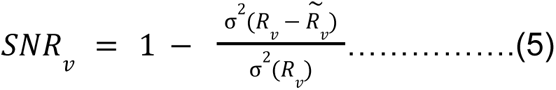

To estimate the signal-to-noise ratio (SNR) of each voxel, we used the test story that was presented five different times to each participant and computed the explainable variance across the different responses, *R_v_*. This computation is described in **Equation 5** where 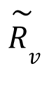 represents the average response across the repeats.

#### Model-based timescale metric T_v_

To estimate the processing timescale *T_v_* of each individual voxel *v* in each participant as per **Equation 2**, we first estimated the contribution of each MT-RNN Layer 2 unit towards modeling *v*’s activity. Since this layer had 1150 multi-timescale features and we used an FIR model with 4 delays to approximate HRFs, MT-EM had to learn a set of 1150 * 4 = 4600 linear weights. To compute the unit contributions, we averaged the weights across the 4 delays and obtained a 1150-D vector β_*v*_. In our prior work^20^, we had estimated β_*v*_ by simply averaging the weights across the FIR delays. While this does not impact long timescale features, high frequency features can be very different across consecutive TRs. Averaging can thus make their contribution tend to 0 and lead to an overemphasis on long timescales. Taking the absolute value of each weight before averaging across delays corrects for this. Additionally, although prior studies have found different HRFs across cortical regions during language processing^90^, the *T_v_* estimates made using weights of each individual FIR delay were highly correlated with each other (whole cortex-µ(*r*) = 0. 82; σ(*r*) = 0. 04; significantly predicted voxels-µ(*r*) = 0. 66; σ(*r*) = 0. 10; *N* = 7). The individual delay estimates were also well correlated with estimates from the average absolute value of weights across all delays (µ(*r*) = 0. 92; σ(*r*) = 0. 02 for whole cortex & µ(*r*) = 0. 85; σ(*r*) = 0. 06 for voxels significantly by MT-EM; *N* = 7). This suggests that there is little difference in HRF shape across cortical voxels based on their timescale.

There are multiple ways we can interpret this model-based timescale metric. Firstly, we can convert this estimate into seconds by multiplying it with the average duration of each word in the stimulus set (0.32 seconds). This gives us a spectrum of timescales from around 0.8 to over 4 seconds. Secondly, this metric can be interpreted as the average exponential decay constant of the MT-RNN units weighted by their contribution towards predicting the voxel’s activity (**Equation 2**). To compare it with prior studies that estimate the exponential decay constant ofindividual neurons, we can compute the half-life, *HL_v_*, of each voxel as *T_v_ * ln(2)* and find the amount of time it takes to decay by ∼99%, ie., approximately 7 * *HL_v_*. This gives us a spectrum of timescales from around 4 to 20 seconds. Lastly, since these estimates are based on MT-RNN layer 2 units that receive input from layer 1 we can also shift the *T_u_* estimates in **Equation 2** by the average timescale of layer 1. Since layer 1 units have *T_u_ E [3, 4]*, the average is 3.5. This would give us a spectrum of timescales from 7.61 to 11.21 words. Although, it is important to note that this is an approximation and each layer 2 unit may not operate exactly at *T_u_ + 3. 5* (see **MT-RNN** details above).

### Framework for *in silico* experiments

To evaluate computational models of language processing, it is not enough to test their prediction performance on held-out stimuli. We also need to determine if these models can reproduce observed brain function under different experimental conditions^91,92^. Here we propose an *in silico* experimentation framework to evaluate this. We demonstrate its use by evaluating the multi-timescale RNN’s ability to capture known temporal context effects in the human cerebral cortex. These effects included the response of different brain regions to scrambled stories^14^, stories in which some words have been replaced by their synonyms^24^, and the rate at which different regions constructed or forgot contextual information^25^.

In this evaluation framework, we first tested how different timescale units in the neural network itself responded to the temporal manipulations. First, we manipulated the stimuli as prescribed in each study and extracted the multi-timescale features of each word in the story from MT-RNN layer 2. Then we adapted the methods used in the original studies to evaluate the effect of the manipulation on each individual MT-RNN unit *u*. Finally, we analyzed if the behavior of *u\* matched its assigned timescale *T_u_*.

Having evaluated the behavior of MT-RNN, we used the learned MT-EM of each voxel to simulate its response to different experimental conditions. We adapted metrics from the original studies to evaluate how the voxel behaved *in silico* and compared these metrics against the model-based estimate of the voxel’s timescale, *T_v_*. We reported experimental results for each voxel. For analyses that involved testing the statistical significance of a correlation, we performed permutation tests. Here we permute one of the dependent variables and recompute the correlation, repeating this for between 1000 and 10000 iterations. We then report the 2-tailed p-value of the permutation test.

To produce group-averaged data from the *in silico* experiments, we followed the procedure reported in the section above to project each subject into a common anatomical space. We then conducted prevalence tests to estimate the statistical significance of the effect at the group-level, again following the procedure described above.

### Temporal scrambling experiment

In Lerner et al. 2011^14^, participants listened to one story from *The Moth Radio Hour* while their fMRI BOLD responses were recorded. The story was either played in its original form, played backwards or played with either its words, sentences or paragraphs scrambled. To estimate the processing timescale of different brain regions with this experiment, the authors first registered all participants into a common-space. Then for each voxel in the common space they computed the inter-subject correlation (ISC) for each stimulus presentation. Finally, the timescale was estimated by determining the conditions in which the voxel had a statistically significant ISC.

We conducted this experiment *in silico* by scrambling our test story (*wheretheressmoke*) at the word, sentence or paragraph level. The main analysis was done on an approximate split, where each story was divided into non-overlapping continuous chunks of a similar temporal duration as in the original study (9 words per sentence & 55 words per paragraph). Alternatively, we manually segmented sentences and paragraphs. We observed little differences in the *in silico* results for the fixed chunks (**Figure 2**) vs. manually-annotated segments (**Supp. Figure S5**).

Since the *in silico* framework does not require new data collection, we could produce many different scrambled versions of the story and repeat the experiment. We reported results across 100 different scramble trials per condition (word, sentence or paragraph). After scrambling the stimuli, we passed each version through MT-RNN and extracted new multi-timescales features for every word from layer 2. Then we used the learned MT-EM to predict the response of each cortical voxel to each scrambled version.

#### Metrics for temporal scrambling experiment

To estimate the effect of scrambling on each voxel, we either (1) compared the variance of its MT-EM predicted response to the intact story and a scrambled version, or (2) re-ordered the scrambled responses to match the original word order and then computed the correlation between the (re-ordered) scrambled and intact responses.

To measure the statistical significance of the response variance metric, we computed the number of scramble trials (out of 100) that had a higher variance than the intact response. We did FDR correction^31^ on these p-values to determine which voxels were significantly affected by the scrambling condition (*q < 0. 05*). These results are reported in **Figure 2**.

For the correlation-based metric, we computed z-scores that described how far the correlation was from 1 (perfect correlation). The standard deviation for the z-score was estimated using the number of samples in each response vector, i.e., the number of TRs. This estimate thus ignored any autocorrelation between TRs and was constant for all voxels. To compute a p-value from the z-scores, we measured the number of scrambling trials for which the z-score was below a specified threshold (-1.65). After this, FDR correction^31^ was applied to determine which voxels were significantly affected by the scrambling condition (*q < 0. 05*). These results are reported in **Figure S4**.

To group voxels into different timescale buckets (*sublexical, word-level, sentence-level* or *paragraph-level & beyond*), we used a cascading procedure. First, all voxels that were significantly predicted under MT-EM were assigned to the *paragraph-level & beyond* bucket. Then voxels whose responses were significant for sentence-level scrambling were assigned to the *sentence-level* bucket. Then voxels whose responses were significant for word-level scrambling were assigned to the *word-level* bucket. Finally, all voxels that were significantly predicted under the sublexical EM with *r > 0. 4* were assigned to the *sublexical* bucket. Overall, we found that the response variance-based timescale estimates closely matched the temporal hierarchy reported in prior studies. In contrast, the correlation-based metric overestimated word-level areas and it was hard to specify a reasonable threshold for the z-scores. We thus report response variance-based estimates in the main text.

We followed the procedure described in an earlier section to project each participant’s timescale metric into fsaverage space, perform significance tests and estimate timescale buckets. To combine data across participants, we computed the arithmetic mode of timescale buckets assigned to each fsaverage voxel. For the *sublexical* bucket, we ran a prevalence test on the sublexical EM performances across all participants and assigned this timescale bucket to any fsaverage voxel that had a prevalent effect. Although we show the group-averaged flatmap from the original study in **Figure 2F**, we were unable to obtain the statistical significance thresholds from the authors and had to rely on manual statistical thresholds for our *in silico* experiment.

Therefore we cannot quantitatively compare their results with ours.

#### Relationship between inter-subject correlation and response variance

While Lerner et al. used inter-subject correlation (ISC) to study the effect of temporal scrambling on cortical responses, our main *in silico* results use response variance. To understand how these measures are related, we first consider an arbitrary voxel in group space and define its mean response timecourse across participants as normally distributed, 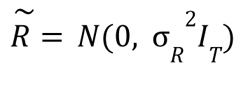 where *T* is the total number of TRs and σ_*R*_^2^ is the response variance. The voxel response for a participant *p* is then given by 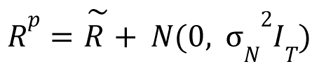 where σ_*N*_^2^ is the noise variance. Since we cannot directly estimate 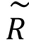, we approximate it as the average response across all participants except *p*, or, 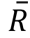. The inter-subject correlation is then given by 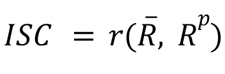. We can see that if σ_*N*_^2^ is held constant, the ISC monotonically increases with the response variance σ_*R*_^2^.

#### Simulating noise for EM predictions to simulate inter-subject correlation

Similar to the cascading procedure we use here, the original study also assigned voxels into timescale buckets by doing a cascading test, but used a different order of assignment. First, the authors assigned all voxels with significant inter-subject correlation (ISC) for the backwards story into the *sublexical* (or, *low-level*) bucket. Then all voxels with significant ISC for word-level scrambling were assigned to the *word-level* bucket. Then voxels with significant ISC for sentence-level scrambling were assigned to the *sentence-level* bucket and finally, all remaining voxels were assigned to the *paragraph-level & beyond* bucket. Aside from the two metrics we discussed above, we also wanted to use the *in silico* framework to more directly replicate ISC and this cascading procedure.

To do this, we first had to simulate and add noise to the MT-EM generated responses. In theory, we can estimate each voxel *v*’s noise variance as *1-D_v_ ^2^* where *D_v_ ^2^* is the coefficient of determination for the EM fit, or, the amount of variance in the voxel’s BOLD responses that can be explained by the MT-RNN features. However, since we do not estimate the regression using ordinary least squares, the generated responses are scaled by the ridge coefficient and 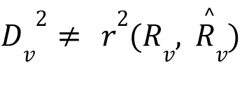. To overcome this, we first z-scored 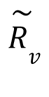 and scaled it by the EM prediction performance, 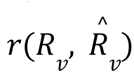, to get 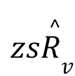. Then, we computed the noise variance as 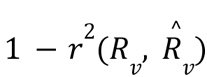 and generated 10 different noise trials of the same length as 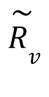 by sampling from a normal distribution. Finally, we simulated ISC for the intact story by adding each noise trial *N_v_* to 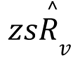 and computing 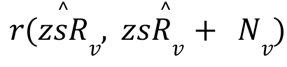. The same procedure was also used to simulate ISC for the different scrambling conditions. We used an exact test to obtain the p-value of each simulated ISC. Following the same cascading procedure as the original study, we assigned voxels into timescale buckets depending on the conditions for which their simulated ISC was significant. The results are shown in **Supp. Figure S4**.

### Narrative divergence experiment

In Yeshurun et al. 2017 ^24^, participants listened to two stories that were matched in content (65.3% identical words), length, and overall duration. The remaining 34.7% of the words were chosen to alter the semantic content of story 1 (S1) and story 2 (S2). For example, "The situation seemed almost *hopeless*, and after *he* spent." in S1 was changed to "The situation seemed almost *too good*, and after *she* spent." As the example illustrates, the word alignment between the stories is not exact--some single words are replaced by two words, some two-word phrases are replaced by single words, etc..

Among other results, they found that in brain regions that integrate over long timescales, the Euclidean distance between the two stories are farther apart than short timescale regions. We repeated this analysis in our in silico replications. To make the distance metric comparable across MT-RNN units, we rescaled each unit’s timecourse to be within [-1, 1] by dividing by its maximum absolute value across both stories. To make the distance metric comparable across different voxels, we rescaled the predicted fMRI responses by the L_2_-norm of its encoding model weights. This is necessary because ridge regression alters the scale of the predicted fMRI outputs, and each voxel has its own ridge regression shrinkage parameter. Finally, we standardized this metric within a participant by re-scaling all distance metrics to fall between 0 and 1. This is done by dividing each voxel metric by the maximum metric within that participant.

We show the group-averaged results following the previously described procedure of projecting individual subject *in silico* data to a common anatomical space.

### Context construction and forgetting experiment

In Chien and Honey 2020, the authors used the same intact and scrambled story dataset as Lerner et al. 2011. However, they compared how different brain regions constructed context over time, or forgot context over time. To do this, they considered the story scrambled in paragraph chunks. For the constructing context analysis, they compared the fMRI responses of a given paragraph chunk when it was preceded by different contexts, or different paragraph chunks. For the forgetting context analysis, they compared the fMRI responses of two different paragraph chunks when they were preceded by the same context. In each region-of-interest (ROI), they computed constructing or forgetting timecourses using inter-participant pattern correlation between the intact and scrambled stories (see **Figure 4A** for a schematic). This procedure involves taking a held-out participant’s pattern of responses and using it to predict the group-averaged pattern of responses, and repeating it while holding out each participant. Here the pattern of interest is across voxels within an ROI, and results in a timecourse for every participant and paragraph chunk. They then generate a single timecourse for each ROI by averaging across participants and chunks.

We pursue a modified version of this analysis that allows us to compute timecourses for each MT-RNN unit. Similar to our in silico replication of Lerner et al. 2011, we used scrambled versions of the test story in the podcast stories dataset. Our main analyses divided the test story into 55-word chunks, which is the average paragraph length. As before, we used 100 different scramble trials. For the MT-RNN features, we reshaped the intact data to form a 3-D matrix of N chunks x 55 words x 1150 MT-RNN units. We excluded the first and last chunks to ensure the same data is used for both the constructing and forgetting analyses, resulting in N = 32 chunks. (This is because the first chunk has no preceding context, and the last chunk has no data following it.) We also reshape the scrambled data to form a 4-D matrix of 100 trials x N chunks x 55 words x 1150 units. Any instance where the scrambled data happens to follow the same order as the intact data (i.e. when the chunks happen to be sequential) is excluded by replacing that scramble trial and chunk with NaN values. Then for every trial and every MT-RNN unit, we constructed a constructing or forgetting timecourse by computing a correlation at each timepoint. The resulting timecourses illustrate whether the pattern of responses across paragraph chunks is similar between the intact and scrambled data. We also repeated this analysis by dividing the MT-RNN units into different timescale bins, and correlating across units within each bin instead of correlating across paragraph chunks. This produced qualitatively similar results.

The analysis for the predicted fMRI data was conducted in a similar fashion. By using a similar analysis as with the MT-RNN, we can maintain single-voxel spatial resolution for each participant and avoid spatial coregistration. First, we determined how many TRs corresponded to each 55-word chunk of the intact story. This varied due to differences in word rate across the story. We then analyzed segments of 9 TRs each, matching the length of data analyzed in Chien and Honey 2020. Any chunks which corresponded to less than 9 TRs were padded with NaN values. We then computed constructing and forgetting context timecourses by correlating the intact data (N chunks x 9 TRs x V voxels) with each trial of the scrambled data (100 trials x N chunks x 9 TRs x V voxels), separately for every voxel. This mirrors the analysis reported for the MT-RNN units, and generates a timecourse for every trial and voxel. We average across trials to show individual voxel results.

The original study authors shared the data containing the constructing and forgetting times for each ROI in the Schaefer atlas^93^. This enabled us to visualize their original effect in **Figure 4D** using pycortex. It also enabled us to show the group-averaged *in silico* results for the same ROI parcellation, following the previously described procedure.

Finally, we also replicated our results by manually segmenting the data at the actual paragraph boundaries for the test story (**Figure S7**).

### Model-free timescale metric based on signal coherence

In order to estimate voxel timescales without relying on the MT-EM, we used a spectral approach. The model-free timescale *T_mf_* of each voxel depends on the spectral coherence between its "true" and recorded BOLD response, i.e., the extent to which the "true" response can be predicted from the recorded response at a particular frequency. This measure can also be thought of as a normalized signal-to-noise measure in the frequency domain. If *S(w)* is the power of the voxel’s true response at some frequency *w* and *N(w)* is the same measure but for noise, then the coherence is calculated as 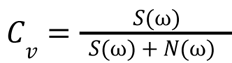. The ‘true’ response was estimated by averaging the voxel’s response across repeats of the test story, and *S(w)* was obtained by taking a periodogram of this averaged voxel response with Welch’s method (64 timepoints per segment; 32 timepoint overlap). The noise was estimated using a jackknife approach. We took a voxel response to one repeat of the test story, subtracted the mean response across all other repeats, and did the same for all other repeats of the test story and aggregated them. The noise power *N(w)* was obtained by taking a periodogram of the jackknifed noise estimate.

In order to estimate *T_mf_*, we performed a weighted average of frequencies for each voxel where the weights for each frequency was the coherence at that frequency. Because coherence for voxels tended to be unimodal across frequencies, this weighted average was close to where coherence was high for each voxel. Finally *T_mf_* is equal to the reciprocal of this weighted average. This inversion means that low values of *T_mf_* correspond to short timescales.

### Probing the timescale of linguistic features in MT-RNN

We built linear classifiers to probe different types of linguistic features in MT-RNN. The classifiers were fit either on the full 1150-D MT-RNN feature space or on a subset of features based on their timescales. For each feature space and task, a classifier was fit in two stages. First, we identified which hyperparameters maximized each classifier’s performance on a validation set. We did grid search on four different learning rates, four learning rate decay constants and three standard deviation values for normally-distributed weight initialization.

Using the best hyperparameters, we then reported performance on a held-out test set with a bootstrapping procedure. Examples were randomly sampled from the test set with replacement 1,000 different times and the mean performance across all runs was reported. We used the 2.5th & 97.5th percentile of the bootstrap distribution to report 95% confidence intervals on classifier performance. For both validation and testing, performance was averaged across two random seeds.

Below we describe the datasets used to probe each linguistic feature.

#### Part-of-speech and Named Entity type

We used the OntoNotes v5.0 dataset^94^ for this task. In this dataset, every word in the document is annotated with its part-of-speech (49 PoS classes in total). Additionally, named entities are divided into 36 classes such as person, location etc. We passed each document through MT-RNN to extract a representation for every token in the document. Then, we trained linear classifiers from the token representation to the annotated part-of-speech or named entity class. We followed the training, validation and test splits specified in the dataset. More details on each split can be found in **Table 1**.

**Table 1:** Ontonotes Dataset for probing part-of-speech and named-entity-recognition from MT-RNN features. We report mean and standard deviation statistics.

|  | TRAINING | VALIDATION | TESTING |
| --- | --- | --- | --- |
| Number of documents | 1933 | 222 | 222 |
| Sentence length | 17.54 ± 12.91 | 16.98 ± 12.88 | 17.89 ± 13.06 |
| Sentence per document | 36.79 ± 67.48 | 43.26 ± 90.40 | 42.70 ± 85.33 |
| Words per document | 645.25 ± 967.34 | 734.70 ± 1184.27 | 763.87 ± 1289.25 |
| Part-of-speech examples | 1247268 | 163104 | 169579 |
| Named-entity-recognition examples | 144308 | 20023 | 20913 |

#### Sentence entailment

We used the Stanford SNLI corpus^95^ for this task. In this task, a classifier is presented with 2 sentences and it has to pick one of 3 possibilities: (1) sentence 1 entails sentence 2, (2) the sentences are in contradiction, (3) they are neutral with respect to each other. To train linear classifiers for this task, we first concatenated the 2 sentences and passed them through the MT-RNN. We extracted the hidden state of layer 1 at the end of the joint sentence pair and trained linear classifier models to predict one of the 3 classes from this representation. We followed the training, validation and test splits specified in the dataset. More details on each split can be found in **Table 2**.

**Table 2:**
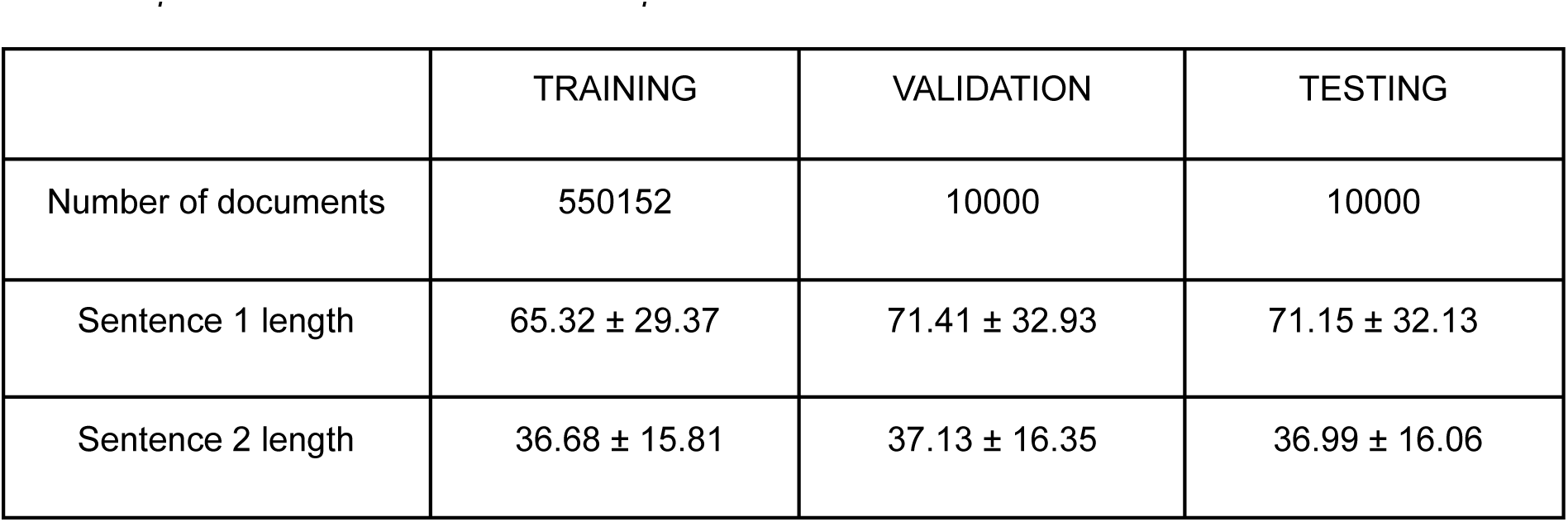
SNLI corpus for probing whether two sentences entail each other, contradict or have no relationship from MT-RNN features. We report mean and standard deviation statistics.

#### Narrative topic

We used the 20Newsgroup dataset for this task (<u>Home Page for 20 Newsgroups Data Set</u>). In this task, each document is annotated with one of 20 topics such as religion, electronics etc. We explored 2 variants to build linear classifier probes. In the first variant, the entire document was passed through the MT-RNN and the hidden state at the end of the document was extracted as a feature on top of which the topic classifier was built. In the second variant, we extracted features for every word in the document and assigned each word the document’s topic label.

We then built a linear classifier from the representation of a word to the topic of the document it belongs to. We followed the train and test split specified in the dataset. About 10% of the training examples were randomly selected to create a validation set. More details on each split can be found in **Table 3**.

**Table 3:** 20Newsgroup dataset for probing narrative topic from MT-RNN features. We report mean and standard deviation statistics.

|  | TRAINING | TESTING |
| --- | --- | --- |
| Number of documents | 11096 | 7370 |
| Document length (in words) | 1027.77 ± 2948.74 | 958.56 ± 2618.16 |

## Supplementary Information

**Supp. Figure S1.**
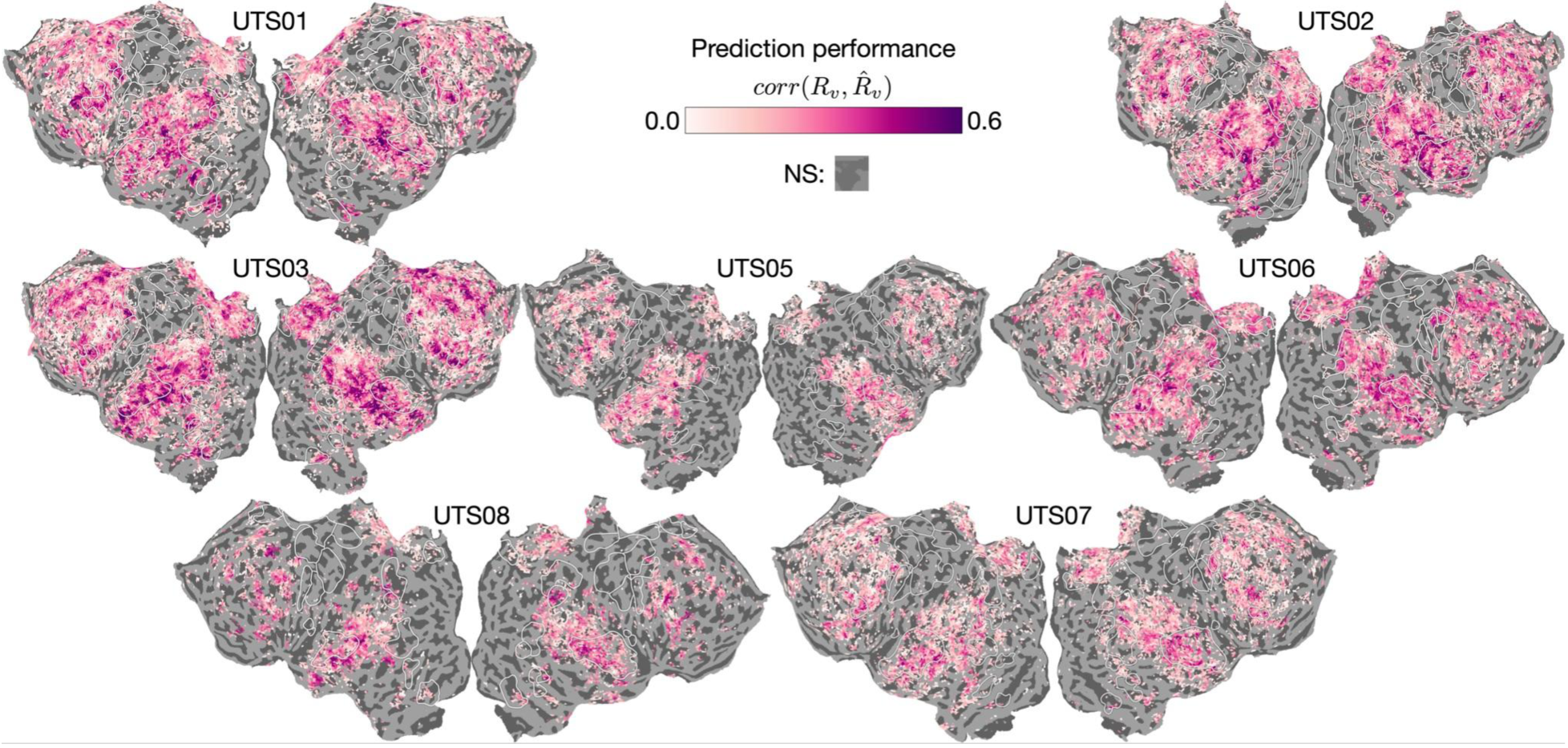
Prediction performance of the multi-timescale encoding models (MT-EM) across all participants. The performance is measured by computing the linear correlation between the true R_v_ and predicted 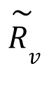 BOLD response for each voxel V on a held-out story that MT-EM has never seen before. All abbreviations follow Figure 1. NS: Not predicted significantly by MT-EM.

**Supp. Figure S2.**
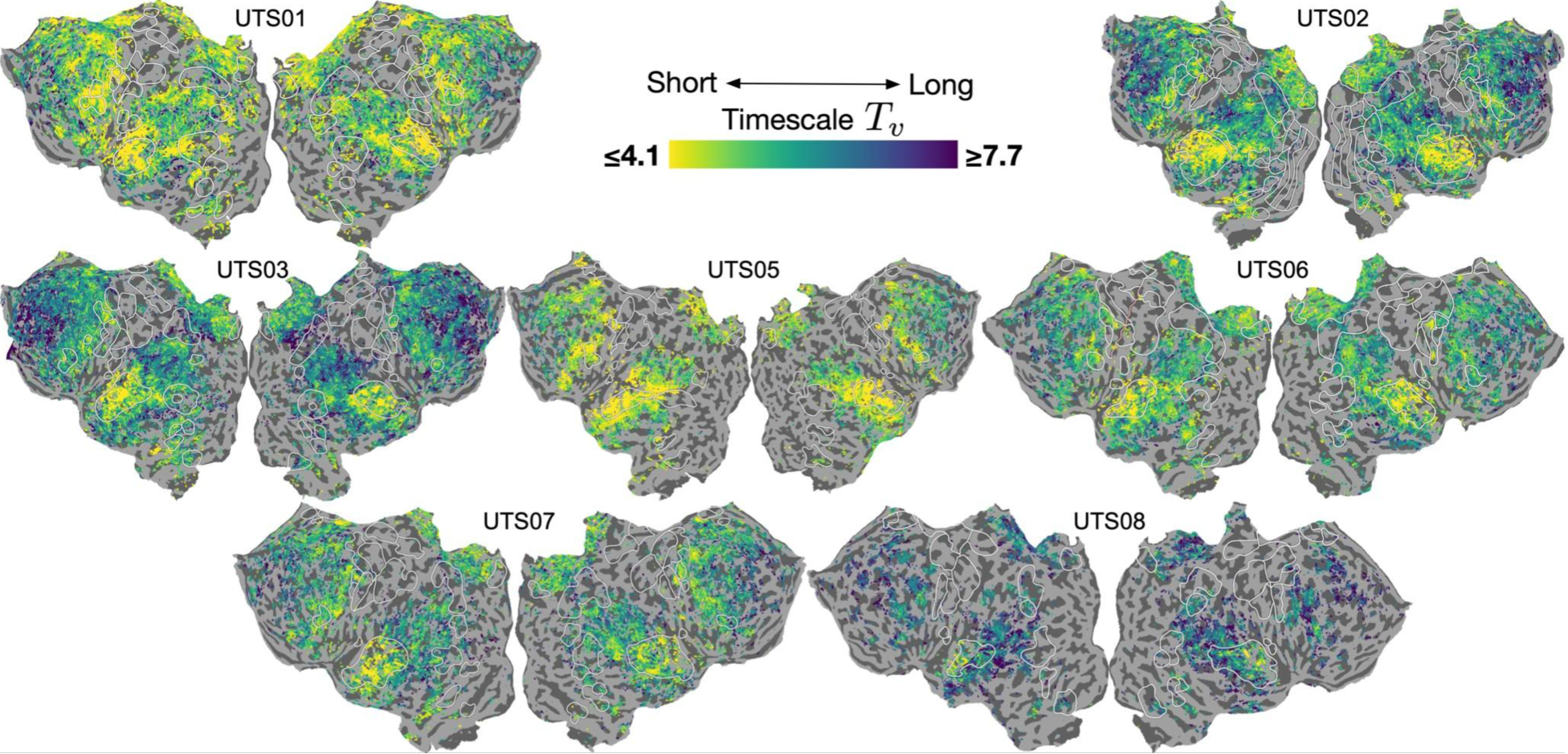
Model-based timescale metric T_v_ for all participants. The metric is computed following ***Equation 2***. All abbreviations follow Figure 1.

**Supp. Figure 3.**
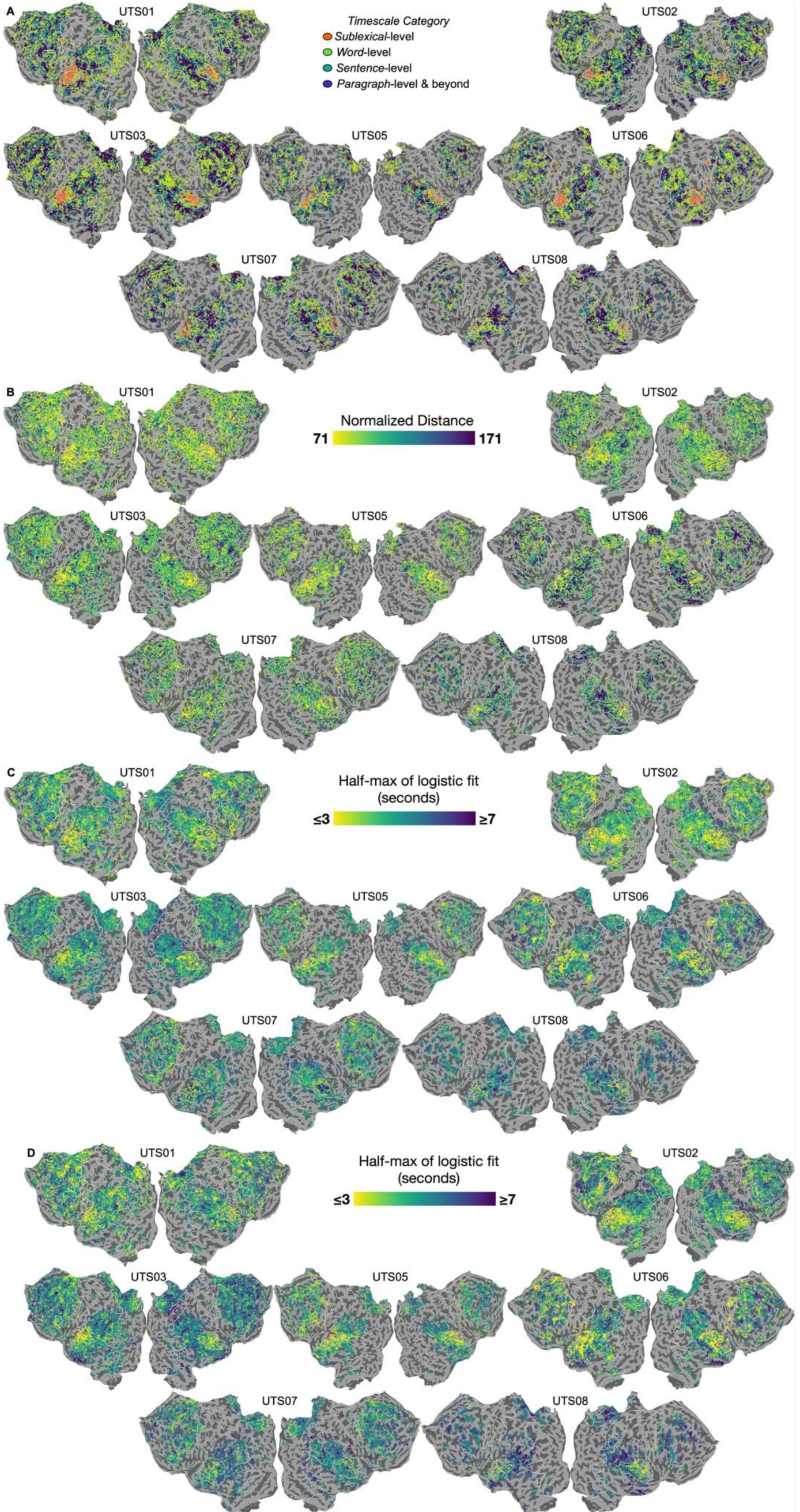
In silico experiment results for all participants. (**A**) Metric is the same as Figure 2E. (**B**) Metric is the same as Figure 3D (**C**) Metric is the same as context construction in Figure 4C. (**D**) Metric is the same as context forgetting in Fig. 4C. All abbreviations follow Figure 1.

**Supp. Figure S4.**
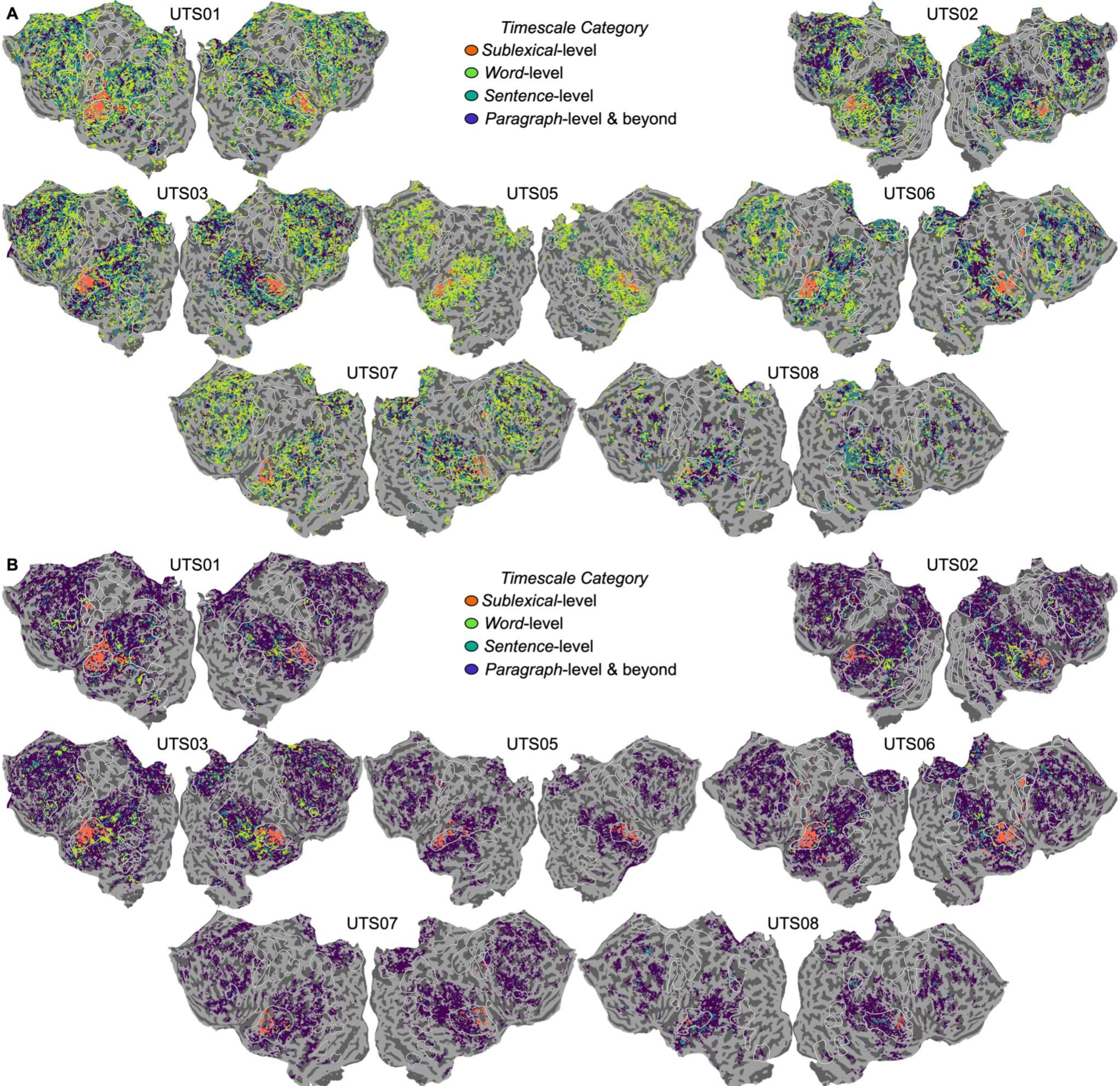
Alternative metrics for the in silico temporal scrambling experiment discussed in Figure 2. (**A**) Correlation-based metric that involves re-ordering the predicted response timecourse for a scrambled story to match the original word order and then computing a linear correlation between the predicted responses for the intact and scrambled versions (see **Methods** for more details). Although the overall trends across cortex match other metrics, we find an overrepresentation of "word-level" regions in most participants. This is similar to results from another replication study^38^ that also used correlation to estimate the effect of temporal scrambling. (**B**) Simulated Inter-Subject Correlation (ISC) obtained by first estimating the noise variance in each voxel of each individual participant and then computing the linear correlation between the original predicted response for a story and noise added on to this response. To classify voxels into four different timescale buckets, the same significance testing and cascading procedure as Lerner et al.^14^ was used (see **Methods** for more details). Although the overall temporal hierarchy matched both prior work^14^ and patterns observed with the other in silico metrics in some participants (Figure 2E & **(A)**), there was an over-representation of long timescales due to a bias in the ISC-based statistical test. In the original study and the simulation here, if a voxel has a significant ISC in a given condition, it implies that the voxel is not affected by the scrambling. In contrast, for our response variance-based timescale estimation, if a voxel has a significant difference in the variance between intact and scrambled responses, it implies that the voxel is affected by the scrambling. Due to the difference in interpretation of statistical significance, the ISC and response variance-based methods are biased in opposite directions. While the ISC-based methods are biased towards the longest timescale bucket (as it does not require the ISC to be significant in any condition), the response variance-based method is biased towards the shortest timescale bucket (as it does not require the response variance to be significantly different from intact in any condition). These biases explain why each metric leads to an over-representation of one timescale category over another and highlights the importance of transparency in estimation procedures. The in silico framework provided the advantage that we could test many different metrics with the same dataset and modeling framework. However, we also observed a strong influence of MT-EM performance in these results-the temporal hierarchy was only evident in participants with very good performance like UTS01-3. Due to the statistical testing bias, around 98.24% of voxels in every participant (a = o. 88; N = 7) with MT-EM performance r < o. 1 were classified into the longest timescale bucket. All abbreviations follow Figure 1.

**Supp. Figure S5.**
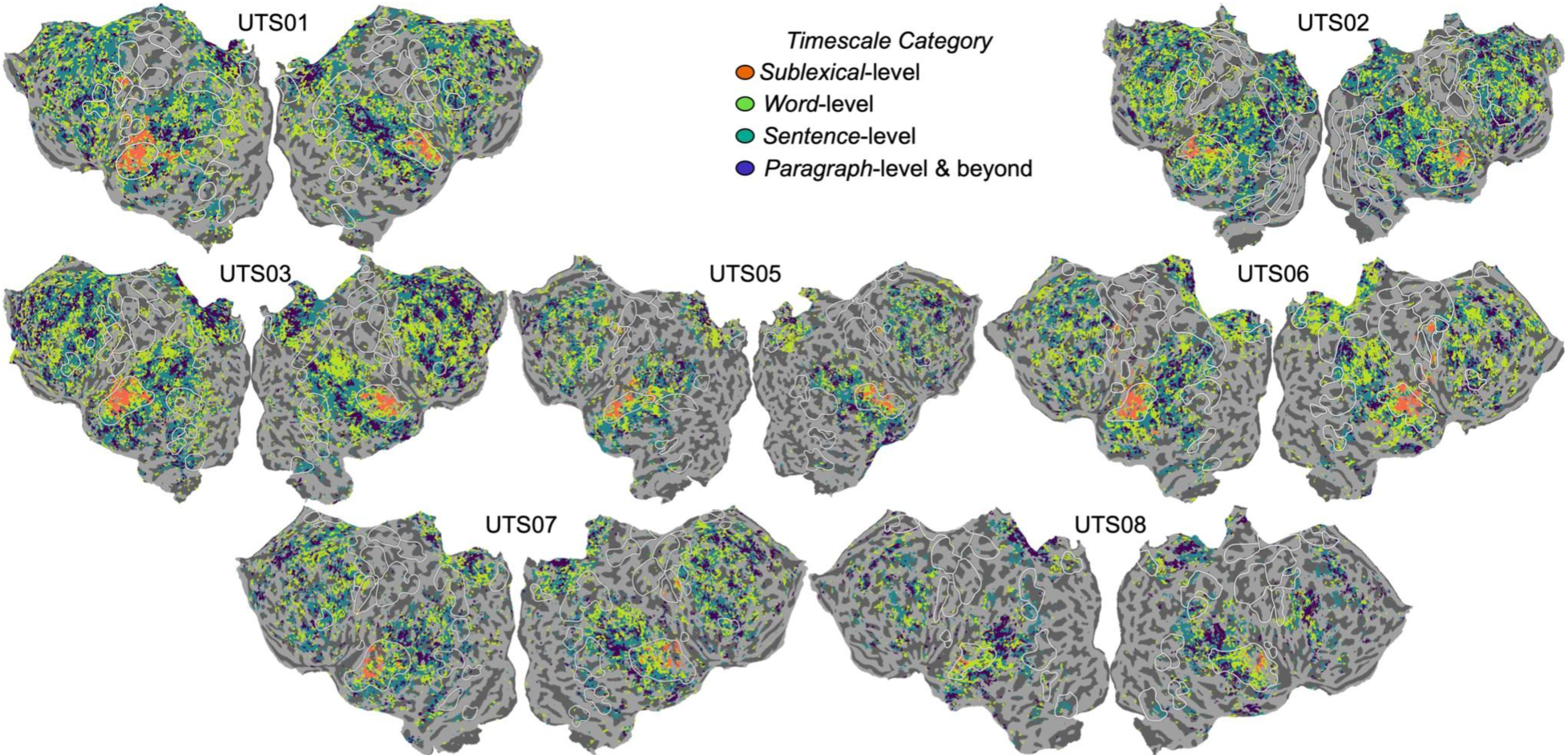
The same as Figure 2D, repeated on the stories manually segmented at the real sentence and paragraph boundaries rather than 9- and 55-word chunks. The pattern of results is the same. All abbreviations follow Figure 1.

**Supp. Figure S6.**
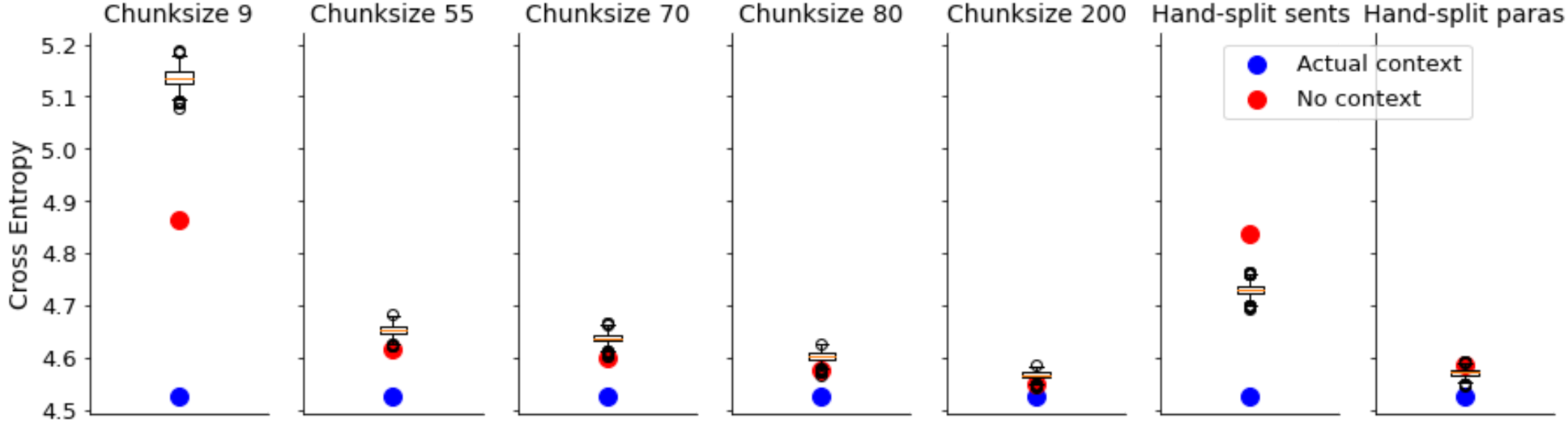
Since MT-EM was unable to replicate the forgetting experiment of Chien & Honey^25^ (Figure 4C-D), we analyzed the forgetting behavior of MT-RNN more closely. First, a test story was divided into chunks of a fixed size (9, 55, 70, 80 or 200 non-overlapping and continuous words) or by manually segmenting into sentences and paragraphs. For each chunk in a story, we then measured MT-RNN’s language modeling performance as the average cross-entropy across all words in the chunk (lower is better) for three different conditions: the chunk was preceded by (1) the correct prior chunk ("correct context") (2) no prior chunks ("no context") or, (3) a randomly-chosen incorrect chunk ("incorrect context"). In every condition, the "correct context" performed better than "no context" or "incorrect context" as expected. Although, for fixed sized chunks, "no context" beat "incorrect context". This suggests that MT-RNN is retaining information in the incorrect prior chunk that leads to worse performance than having no context at all. In manually segmented chunks, "incorrect context" performs better than "no context" suggesting that as long as the grammatical boundaries are intact, MT-RNN can even use incorrect context to perform better at language modeling. Taken together, these two findings show that MT-RNN retains information from prior chunks and does not have a forgetting mechanism like the cerebral cortex.

**Supp. Figure S7.**
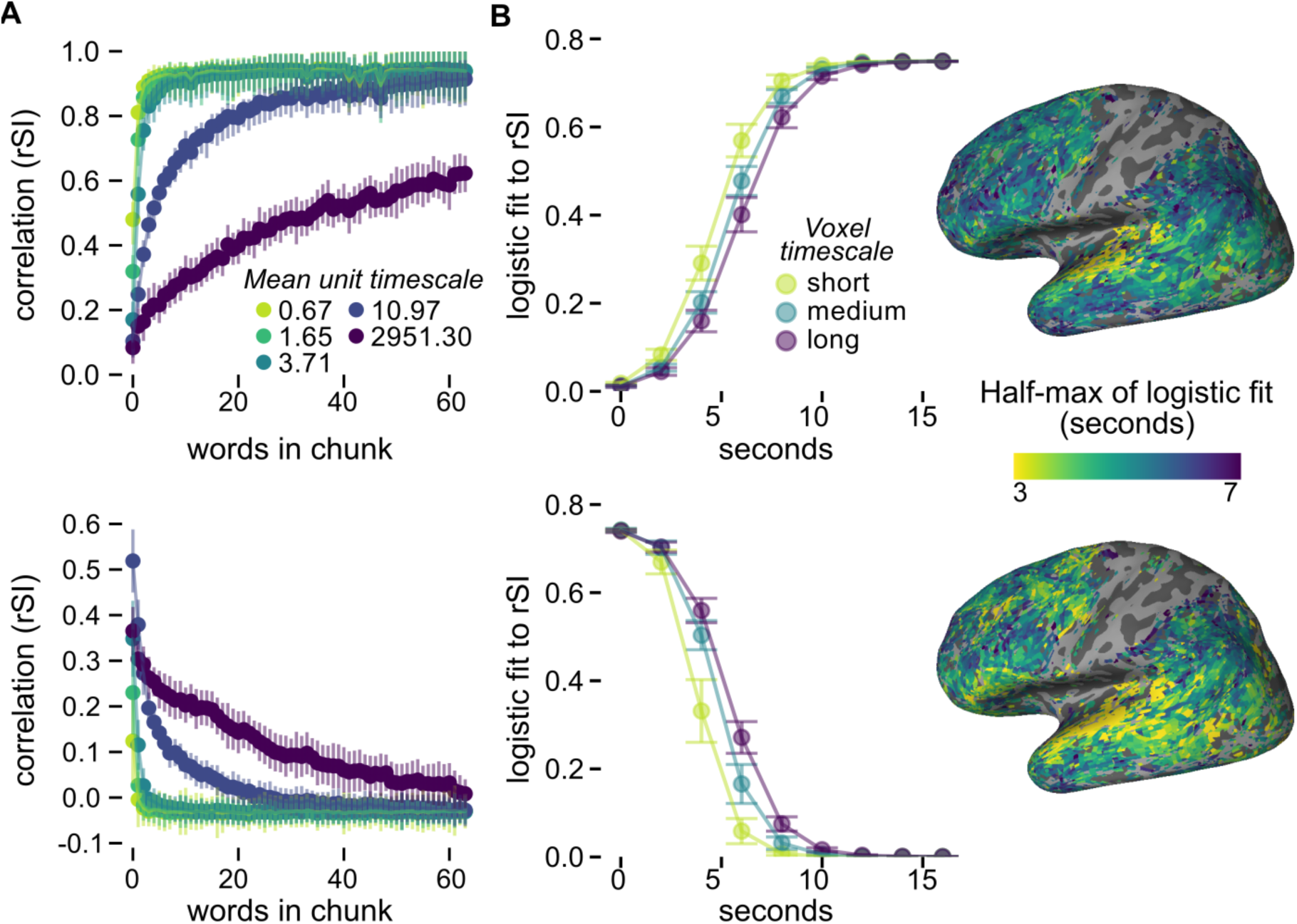
The same as Figure 4, repeated on the stories segmented at the real sentence and paragraph boundaries rather than 9- and 55-word chunks. The pattern of results is the same.

**Supp. Figure S8.**
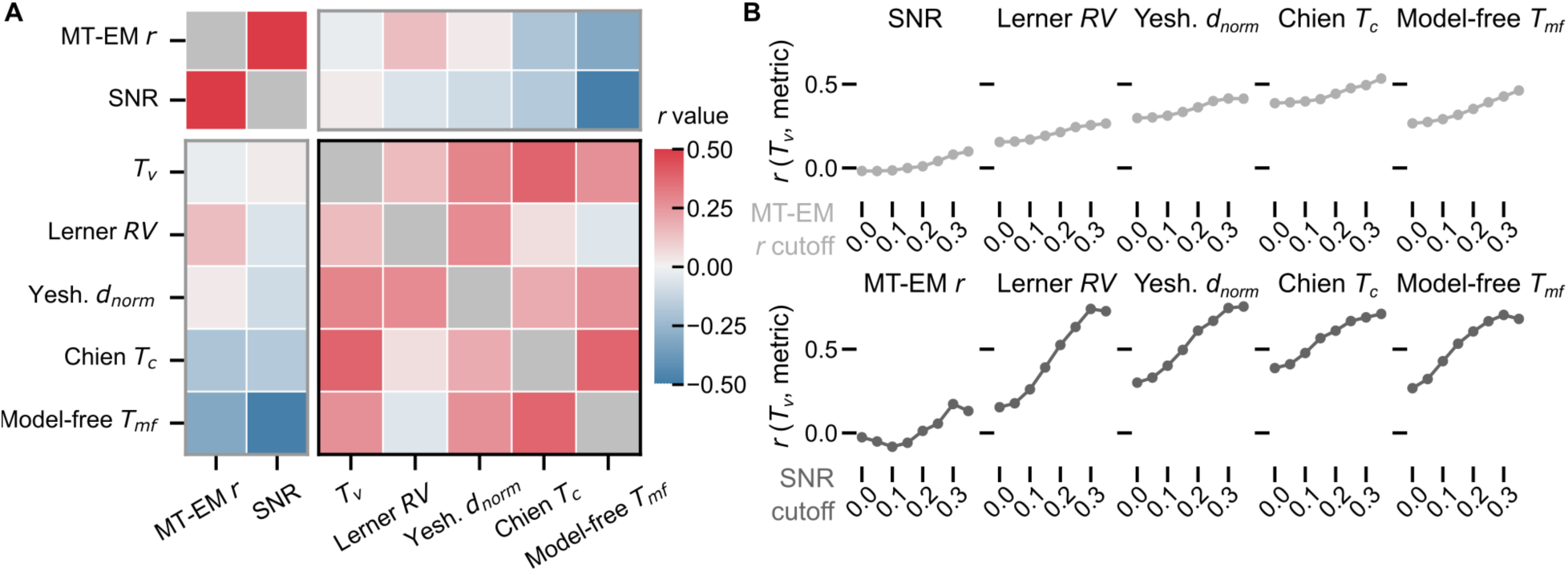
Similar to main text Figure 5, but using metrics projected into the same anatomical space and averaged across subjects. We show the relationship between MT-EM derived timescale T_v_, the in silico experiment metrics that successfully replicate the original reports, the model-free timescale metric T_mf_ based on spectral information, and other metrics related to the fMRI signal. Abbreviations: Lerner RV is the timescale category based on the response variance of the timecourse (see Supplement).Yeshurun d_norm_ is the normalized Euclidean distance between the stories. Chien T is the half-max of the logistic fit for the constructing context condition, while Chien T_f_ is the same for the forgetting context condition. MT-EM r is the measure of encoding model performance, and SNR is the signal-to-noise ratio. (**A**) Cross-correlation matrix of timescale metrics across voxels significantly predicted by the encoding model. (**B**) Correlations re-computed only for voxels that survive a threshold on encoding model performance (top) or SNR (bottom). Similar to Figure 5, this shows that the T_v_ metric more reliably predicts the in silico timescale metrics as signal quality and encoding model performance become more reliable. As the thresholds increase, the correlation between timescale increases. T_v_ and the in silico experiment metrics generally increases.

**Supp. Figure S9.**
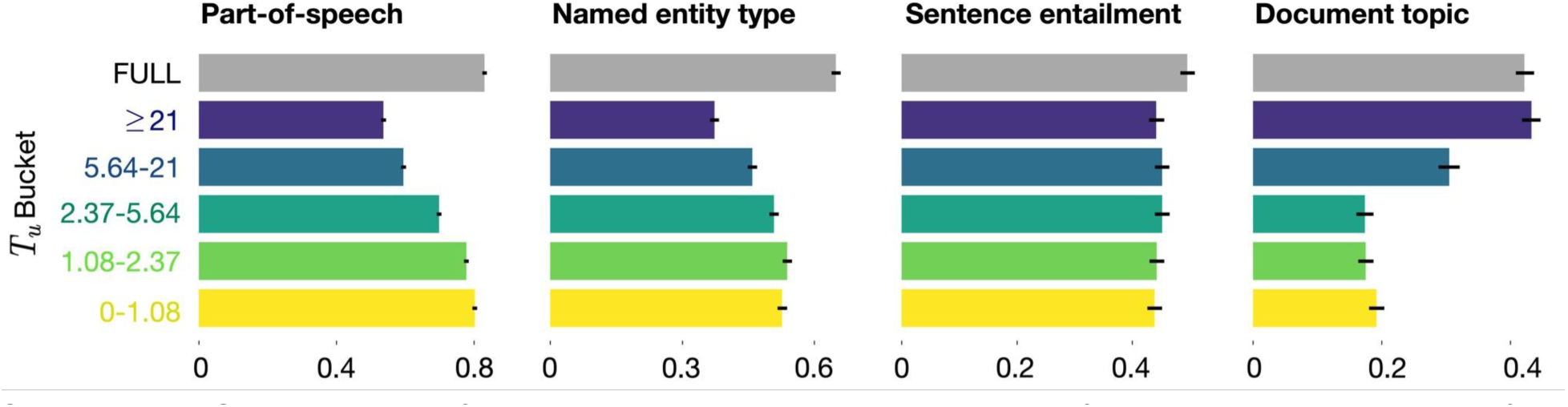
Linear classification results when the 1150-D MT-RNN feature space is divided into five equal splits of 230 units each, as opposed to the linguistically determined splits used in Figure 6. The procedure and results match Figure 6.

