## Supplementary PDF for "A unifying computational account of temporal context effects in language across the human cortex"

### Supplementary Information

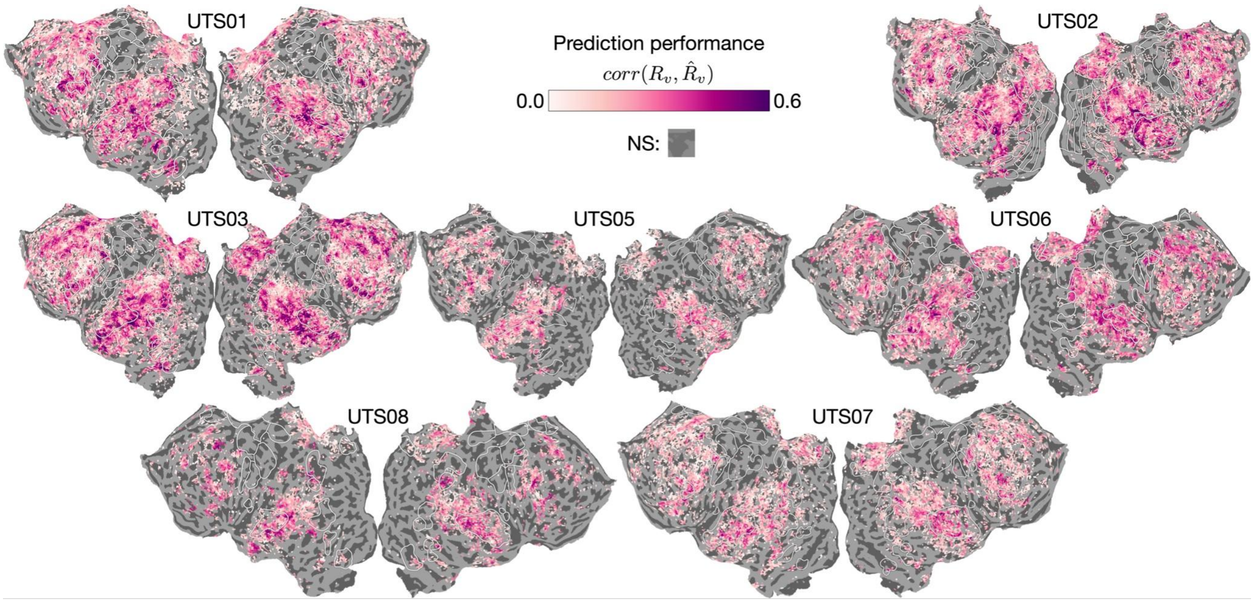


***Supp. Figure S1.*** *Prediction performance of the multi-timescale encoding models (MT-EM) across all participants. The performance is measured by computing the linear correlation between the true* $R_v$ *and predicted* $\hat{R}_v$ *BOLD response for each voxel* $v$ *on a held-out story that MT-EM has never seen before. All abbreviations follow* ***Figure 1****. NS: Not predicted significantly by MT-EM.*

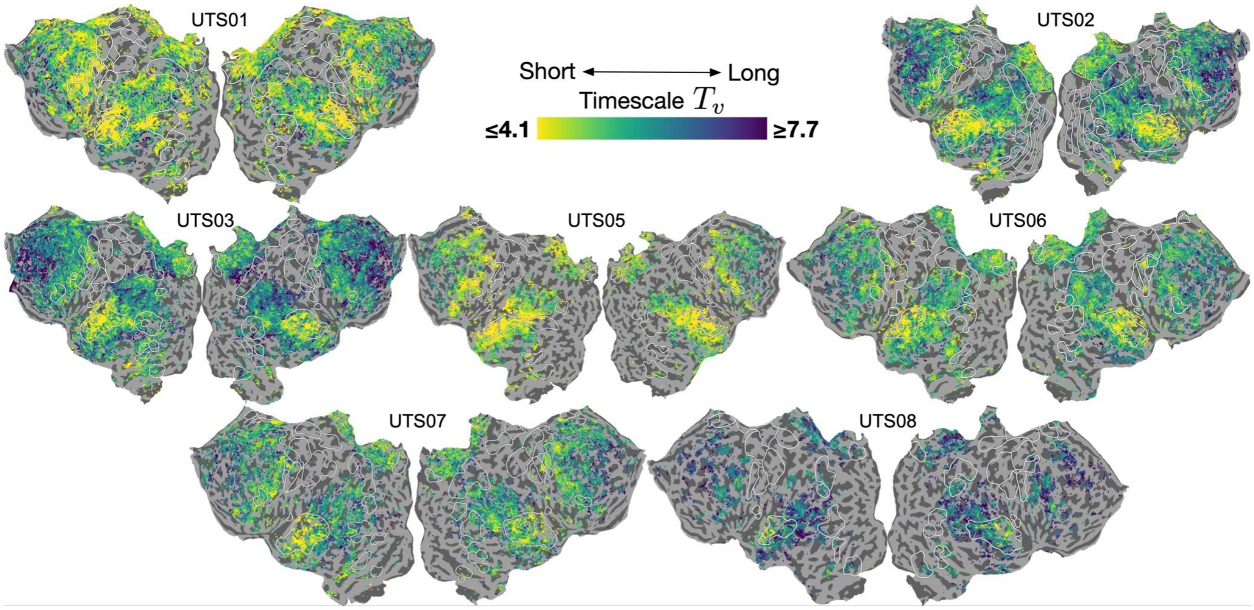


***Supp. Figure S2.*** *Model-based timescale metric $T_v$ for all participants. The metric is computed following* ***Equation 2****. All abbreviations follow* ***Figure 1****.*

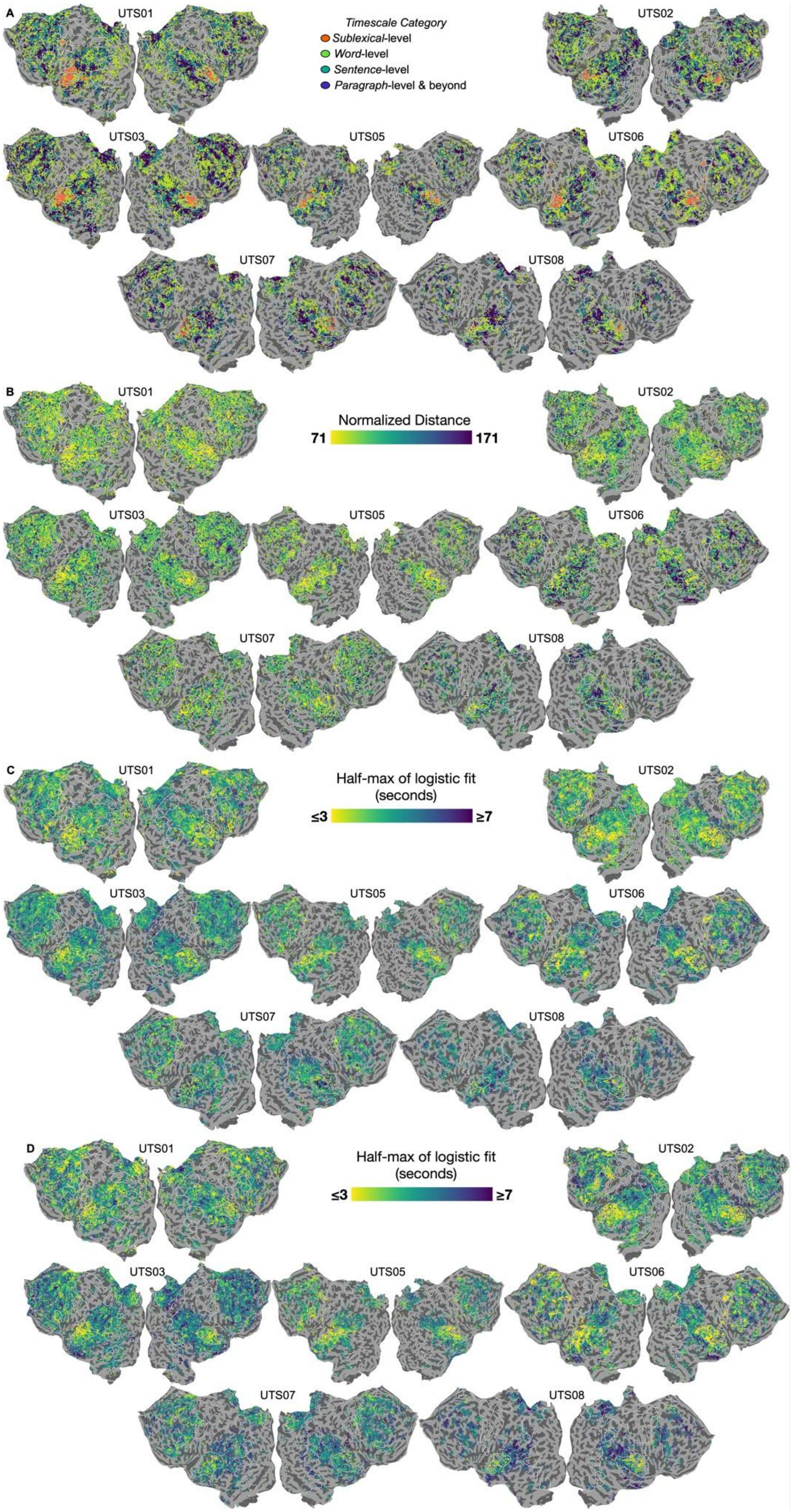


***Supp. Figure 3**. In silico experiment results for all participants. (**A**) Metric is the same as **Figure 2E**. (**B**)*

*Metric is the same as* ***Figure 3D*** *(****C****) Metric is the same as context construction in* ***Figure 4C****. (****D****) Metric is the same as context forgetting in Fig. 4C. All abbreviations follow* ***Figure 1****.*

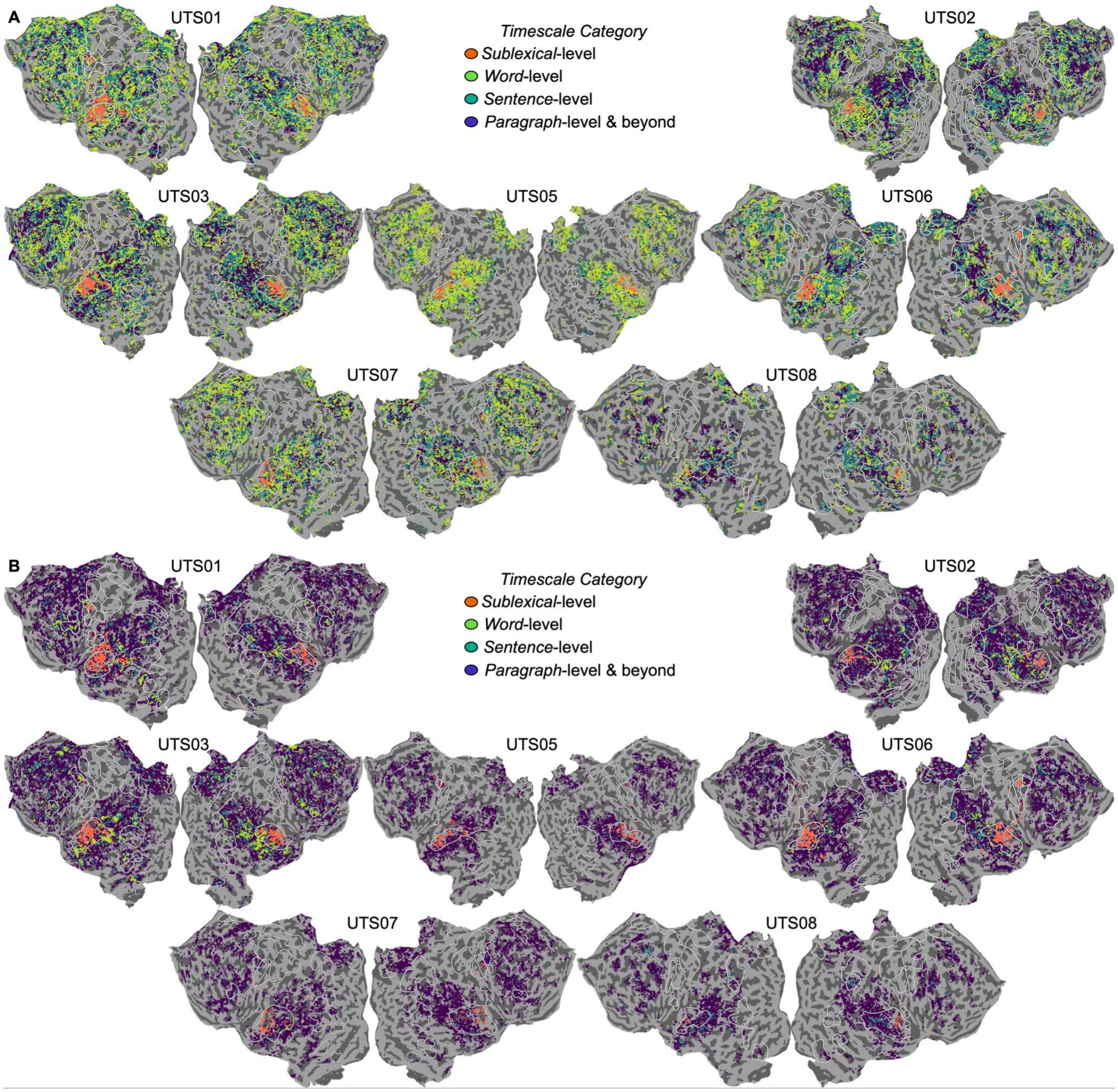


***Supp. Figure S4.*** *Alternative metrics for the in silico temporal scrambling experiment discussed in* ***Figure 2****. (****A****) Correlation-based metric that involves re-ordering the predicted response timecourse for a scrambled story to match the original word order and then computing a linear correlation between the predicted responses for the intact and scrambled versions (see* ***Methods*** *for more details). Although the overall trends across cortex match other metrics, we find an overrepresentation of "word-level" regions in most participants. This is similar to results from another replication study[38] that also used correlation to estimate the effect of temporal scrambling. (****B****) Simulated Inter-Subject Correlation (ISC) obtained by first estimating the noise variance in each voxel of each individual participant and then computing the linear correlation between the original predicted response for a story and noise added on to this response. To classify voxels into four different timescale buckets, the same significance testing and cascading procedure as Lerner et al.[14] was used (see* ***Methods*** *for more details). Although the overall temporal hierarchy matched both prior work[14] and patterns observed with the other in silico metrics in some participants (****Figure 2E*** *&* ***(A)****), there was an over-representation of long timescales due to a bias in the*

*ISC-based statistical test. In the original study and the simulation here, if a voxel has a significant ISC in a given condition, it implies that the voxel is not affected by the scrambling. In contrast, for our response variance-based timescale estimation, if a voxel has a significant difference in the variance between intact and scrambled responses, it implies that the voxel is affected by the scrambling. Due to the difference in interpretation of statistical significance, the ISC and response variance-based methods are biased in opposite directions. While the ISC-based methods are biased towards the longest timescale bucket (as it does not require the ISC to be significant in any condition), the response variance-based method is biased towards the shortest timescale bucket (as it does not require the response variance to be significantly different from intact in any condition). These biases explain why each metric leads to an over-representation of one timescale category over another and highlights the importance of transparency in estimation procedures. The in silico framework provided the advantage that we could test many different metrics with the same dataset and modeling framework. However, we also observed a strong influence of MT-EM performance in these results— the temporal hierarchy was only evident in participants with very good performance like UTS01-3 . Due to the statistical testing bias, around 98.24% of voxels in every participant ($\sigma = 0.88$; $N = 7$) with MT-EM performance $r < 0.1$ were classified into the longest timescale bucket. All abbreviations follow* ***Figure 1***.

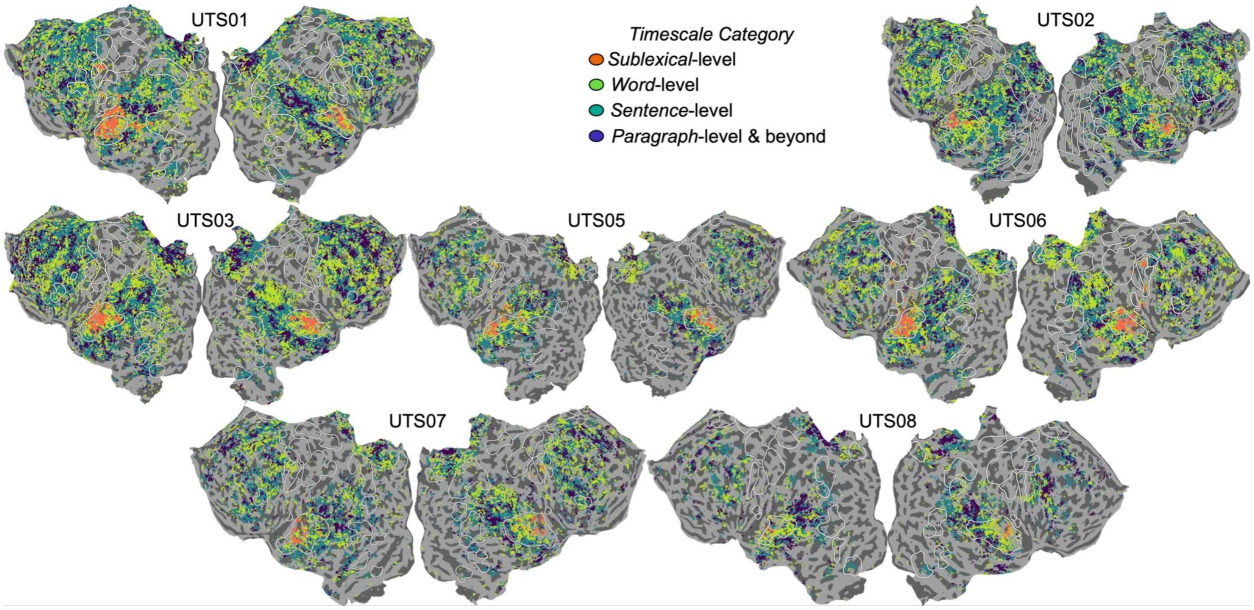


***Supp. Figure S5.*** *The same as* ***Figure 2D****, repeated on the stories manually segmented at the real sentence and paragraph boundaries rather than 9- and 55-word chunks. The pattern of results is the same. All abbreviations follow* ***Figure 1***.

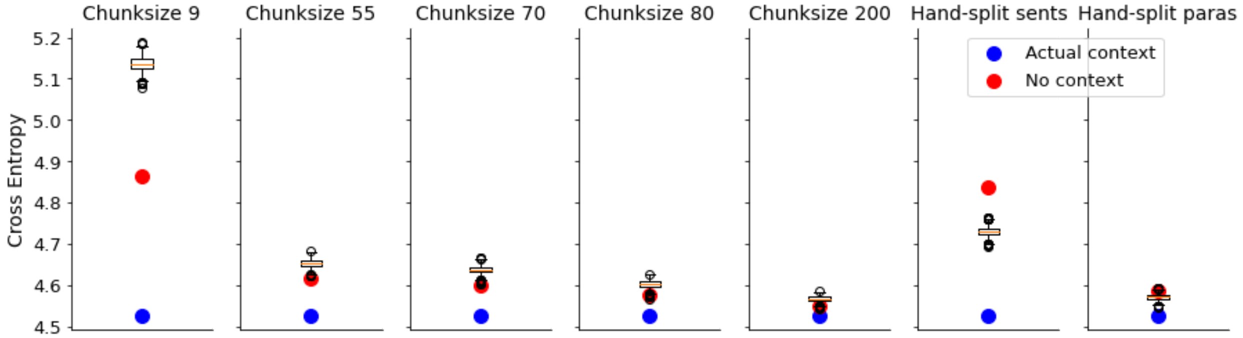


***Supp. Figure S6.*** *Since MT-EM was unable to replicate the forgetting experiment of Chien & Honey*[25] *(****Figure 4C-D****), we analyzed the forgetting behavior of MT-RNN more closely. First, a test story was*

*divided into chunks of a fixed size (9, 55, 70, 80 or 200 non-overlapping and continuous words) or by manually segmenting into sentences and paragraphs. For each chunk in a story, we then measured MT-RNN's language modeling performance as the average cross-entropy across all words in the chunk (lower is better) for three different conditions: the chunk was preceded by (1) the correct prior chunk ("correct context") (2) no prior chunks ("no context") or, (3) a randomly-chosen incorrect chunk ("incorrect context"). In every condition, the "correct context" performed better than "no context" or "incorrect context" as expected. Although, for fixed sized chunks, "no context" beat "incorrect context". This suggests that MT-RNN is retaining information in the incorrect prior chunk that leads to worse performance than having no context at all. In manually segmented chunks, "incorrect context" performs better than "no context" suggesting that as long as the grammatical boundaries are intact, MT-RNN can even use incorrect context to perform better at language modeling. Taken together, these two findings show that MT-RNN retains information from prior chunks and does not have a forgetting mechanism like the cerebral cortex.*

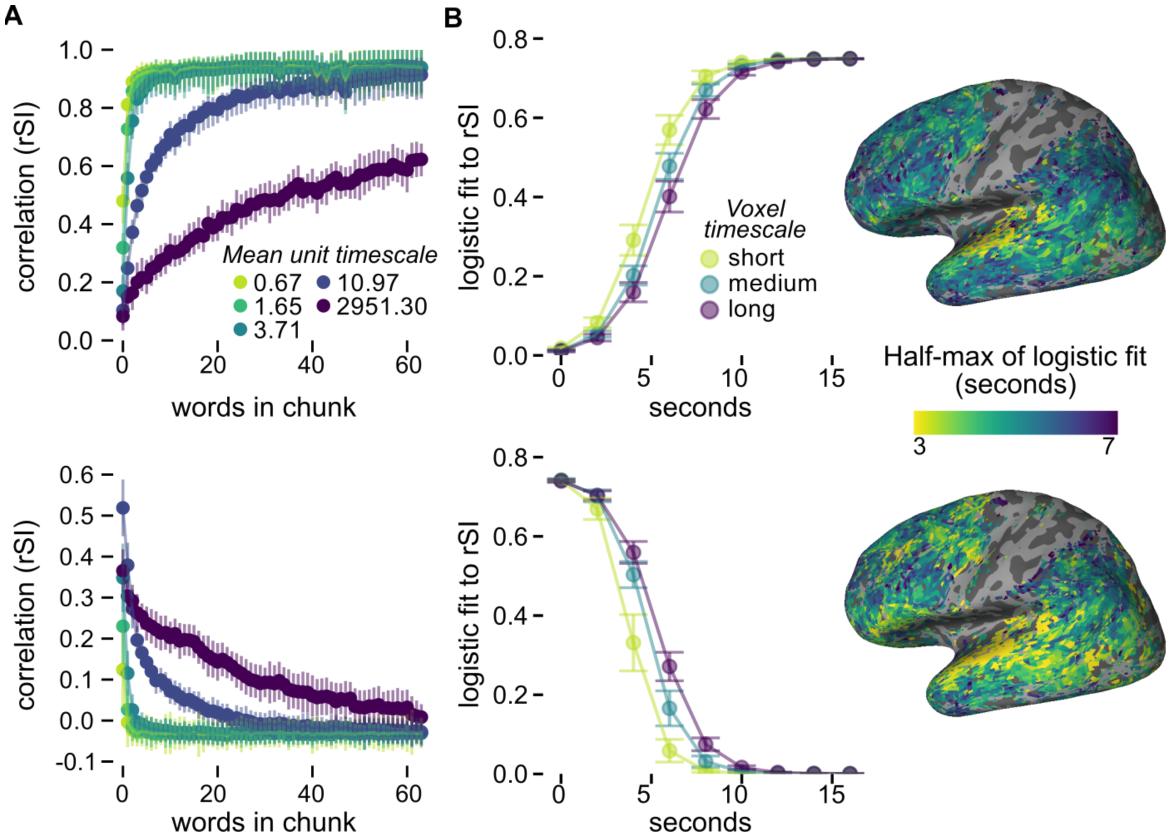


***Supp. Figure S7.*** *The same as* ***Figure 4****, repeated on the stories segmented at the real sentence and paragraph boundaries rather than 9- and 55-word chunks. The pattern of results is the same.*

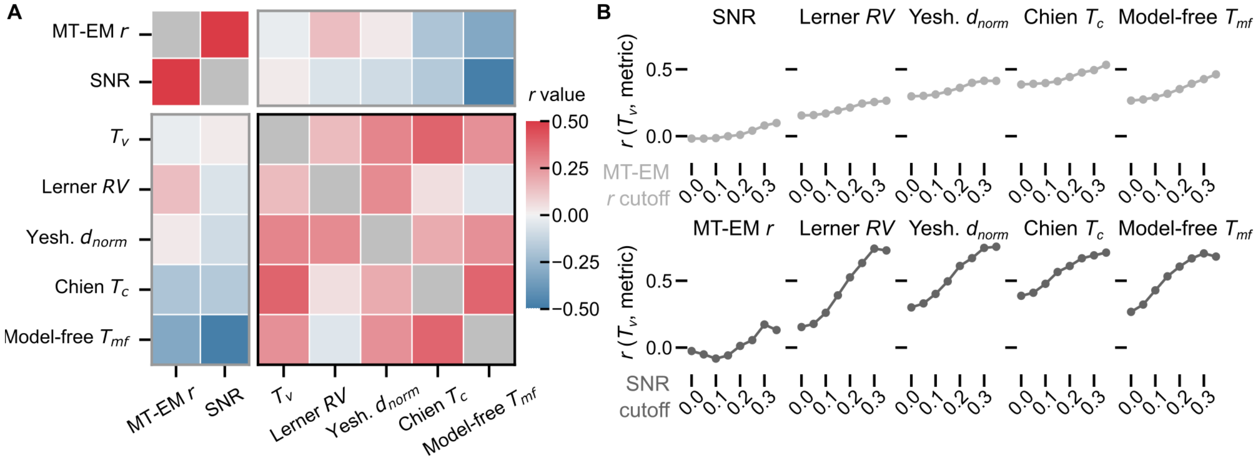


***Supp. Figure S8.*** *Similar to main text* ***Figure 5****, but using metrics projected into the same anatomical space and averaged across subjects. We show the relationship between MT-EM derived timescale $T_v$, the in silico experiment metrics that successfully replicate the original reports, the model-free timescale metric $T_{mf}$ based on spectral information, and other metrics related to the fMRI signal. Abbreviations: Lerner $RV$ is the timescale category based on the response variance of the timecourse (see Supplement).Yeshurun $d_{norm}$ is the normalized Euclidean distance between the stories. Chien $T_c$ is the half-max of the logistic fit for the constructing context condition, while Chien $T_f$ is the same for the forgetting context condition. MT-EM r is the measure of encoding model performance, and SNR is the signal-to-noise ratio. (****A****) Cross-correlation matrix of timescale metrics across voxels significantly predicted by the encoding model. (****B****) Correlations re-computed only for voxels that survive a threshold on encoding model performance (top) or SNR (bottom). Similar to* ***Figure 5****, this shows that the $T_v$ metric more reliably predicts the in silico timescale metrics as signal quality and encoding model performance become more reliable. As the thresholds increase, the correlation between timescale $T_v$ and the in silico experiment metrics generally increases.*

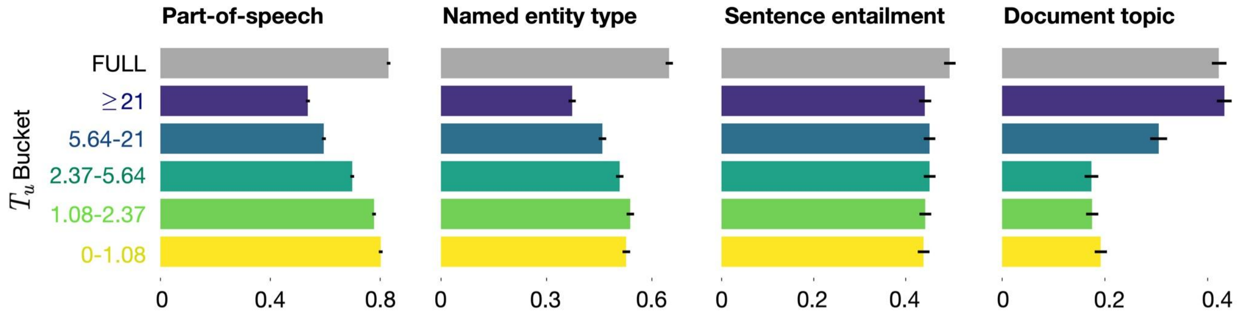


***Supp. Figure S9.*** *Linear classification results when the 1150-D MT-RNN feature space is divided into five equal splits of 230 units each, as opposed to the linguistically determined splits used in* ***Figure 6****. The procedure and results match* ***Figure 6****.*
